# Opposing cell type preferences for binding and replication shape influenza A virus infection in human airways

**DOI:** 10.64898/2026.05.04.722582

**Authors:** Josephine von Kempis, Marie O. Pohl, Elisabeth Gaggioli, Cyrille Niklaus, Alexandra Trkola, Geert-Jan Boons, Robert P. de Vries, Silke Stertz

**Author notes:** Corresponding author: Silke Stertz, Ph.D.

## Abstract

Influenza A viruses (IAVs) pose a persistent threat to human health through seasonal epidemics and zoonotic spillover from avian reservoirs. As respiratory pathogens, they primarily target the airway epithelium. However, it remains unclear how host cell-specific barriers jointly shape viral tropism and replication in primary human airway cultures. Here, we show that avian IAVs can infect ciliated and secretory cells but preferentially bind to ciliated cells, consistent with higher abundance of their receptor α2,3-linked sialic acids, specifically sialyl Lewis X glycans, present on the apical surface of ciliated cells. Replication levels were comparable between secretory and ciliated cells for the avian strains, resulting in an overall preference for ciliated cells. In contrast, human IAVs also preferentially bind to ciliated cells but independently of α2,6-linked sialic acid abundance. Human IAVs replicate more efficiently than avian IAVs due to their ability to utilize human ANP32 proteins, but they also exhibit cell type–specific differences due to ANP32, allowing for higher viral RNA levels in secretory cells. Thus, preferential binding to ciliated cells coupled with enhanced replication in secretory cells equalizes overall infection levels across cell types for human IAVs. Together, our findings highlight the spatiotemporal complexity and interplay of IAV infection dynamics in the airway epithelium and redefine current models of influenza A virus tropism.

## INTRODUCTION

Influenza A viruses (IAVs) cause respiratory tract infections in humans and represent a major challenge to global health and the economy. Beyond the yearly ‘flu’ season (1) responsible for an estimated 1 billion infections, causing 3–5 million cases of severe illness and 290,000–650,000 deaths annually (2) IAVs can pose a pandemic threat (3,4). This arises from the ability of human IAV to undergo genetic reassortment with viruses from their natural avian reservoir, enabling zoonotic spillover and rapid adaptation to new hosts. Understanding the determinants that govern IAV adaptation to human cells is therefore critical for pandemic preparedness.

As the virus enters the human body through the respiratory tract, the airway epithelium is the initial site of viral attachment and infection (5). The airway epithelium is a heterogeneous and highly structured microenvironment, composed of a variety of cell subtypes distributed in a polarized fashion along the basal-apical axis: Basal cells differentiate through intermediate stages to secretory cells, responsible for apical mucus production, and subsequently to ciliated cells, which drive mucosal clearance through ciliary beating (6–9). Additionally, the epithelium also comprises minor cell type populations, such as ionocytes and tuft cells.

For zoonotic infections to occur in the human airway, avian IAV must overcome multiple species-specific barriers. A variety of these barriers have been characterized (10), including differential receptor specificity (11), polymerase incompatibility with human ANP32 proteins (12), lower pH stability of the envelope protein (13,14), and sensitivity to antiviral factors such as Mx (15,16) and BTN3A3 (17). At the same time, the initial interaction of the virus with the heterogeneous airway epithelium raises a fundamental question: which cell subtypes are preferentially infected by different IAV strains, and which host and viral factors impact this process? In 2004, Matrosovich *et al.* reported that human-adapted IAV (hIAV) infect predominantly non-ciliated cells, while avian IAV (aIAV) infect mainly ciliated cells (18). These patterns directly correlated with the distribution of the viral attachment receptors (19–21): While the avian preferred receptors (α2.3-linked sialic acids) were predominantly detected on ciliated cells, sialic acids with an α2.6-linkage, which are the binding receptors of hIAV, were mainly on non-ciliated cells. However, subsequent work has revealed that the infection patterns in the airway epithelium are more complex and less clear-cut than initially suggested. Specifically, both α2.3- and α2.6-linkages have been detected on ciliated and non-ciliated cells (22–24), and hIAV can also infect or at least bind to ciliated cells (23,25). More recent work focused on pandemic 2009 H1N1 strains and reported that its primary targets are secretory cells, while ciliated cells are the secondary targets (26), and ultimately that the ciliated cells are the main driver of hIAV H1N1 viral particle production (24). Together, these findings reopen the fundamental question regarding the IAV cellular tropism and the underlying viral and cellular determinants.

Here, we reveal that human H1N1 and H3N2 IAV infect ciliated and secretory cells at comparable levels in the initial infection round in the human bronchial epithelium. In contrast, avian influenza A strains (H1N1, H3N8, and H11N6) display a clear preference, but not restriction, for ciliated cells. We further show that the avian virus preferentially binds to ciliated cells, driven the by higher abundance of α2.3-linked sialic acids, specifically sialyl Lewis X (sLe^x^) structures, on the apical surface of ciliated cells. We find that human strains also preferentially bind ciliated cells but exhibit higher RNA production in secretory cells due to differential ANP32 expression levels between the cells, resulting in a dual-tropism phenotype. Lastly, we show that the observed deficiency in avian virus replication is not driven by the initial upregulation of interferon-stimulated gene transcription but instead by the limited compatibility of the viral polymerase with human ANP32 proteins.

## RESULTS

### Human and avian influenza A viruses display differential tropism in bronchial human epithelial cells

To model the heterogeneity and complexity of the airway epithelium *in vitro*, we utilized differentiated primary human bronchial epithelium cells (BEpC) grown at air-liquid interface from three independent donors (donors 1-3). The presence of the main bronchial cell types (ciliated, secretory, and basal cells) was determined and quantified using a previously described flow cytometry panel (27) (Fig. 1a). We successfully identified all three cell type populations within each donor (Fig. 1b). To assess the efficiency of the initial infection in BEpC with a human (hIAV) and an avian influenza A virus (aIAV), we used a seasonal human H1N1 IAV from 2019 (A/Hawaii/70/2019) and an avian H1N1 isolated from a duck (A/Duck/Alberta/35/1976). We infected the cultures with three different multiplicities of infection (MOI: 1, 5, and 10) as calculated by titration on MDCK cells and quantified the percentage of infected cells (NP-positive cells) via immunofluorescent microscopy (IF; Fig. 1c, Supp. Fig. 1a). At an MOI of 1, approximately 10-15% of cells were NP-positive for both viruses. The proportion of infected cells increased with higher MOI and reached a plateau at approximately 30% at MOI 5, with no substantial further increase at MOI 10. For flow cytometric detection, infected cells were identified by surface haemagglutinin (HA) staining, which showed insufficient sensitivity at 6 hours post-infection (hpi) but a more robust signal at 8hpi (Supp. Fig.1b). Therefore, we proceeded with flow cytometric readouts at an MOI of 5 and at 8 hpi for both viral infections across all three donors. For subsequent analysis we pooled the findings from donors 1-3. Contrary to previous results (18), both, the avian and the human IAV, infected ciliated and secretory cells (Fig. 1d), whereas infections of basal cells remained minimal. To compare the likelihood of secretory versus ciliated cells becoming infected by a virus while minimizing the effect of between-repeat variability in overall infection levels, we normalized subtype-specific infection rates to the overall percentage of infected cells. We thus yielded a *normalized infection rate*, which revealed that the avian H1N1 strain has a significant preference to infect ciliated over secretory cells, while the human strain did not display a clear bias towards either subtype (Fig. 1e). We further found that dual tropism for both cell types was true for additional H1N1 but also H3N2 human strains and tropism for ciliated cells could be confirmed for additional avian strains (Fig. 1e-f). Lastly, we quantified viral protein levels within the infected cell and found that cells infected with A/Hawaii/70/2019 showed significantly higher viral NP signals than A/Duck/Alberta/35/1976 infected cells (Fig. 1g; Supp. Fig. 1d). These data indicate that differences in viral protein expression efficiency are evident already at early time points post-infection.

**Figure 1.**
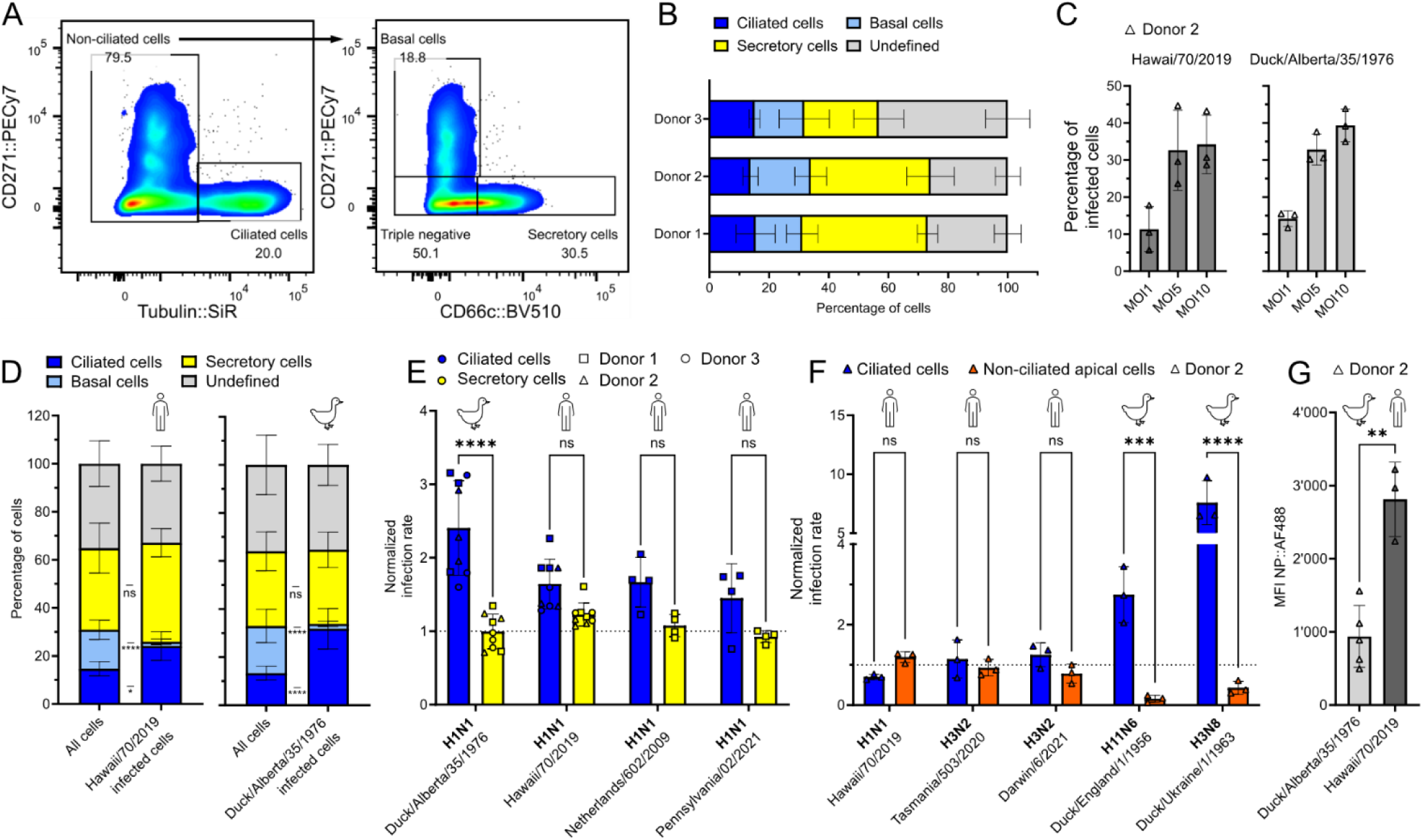
Flow cytometry and microscopy analysis reveal divergent bronchial epithelial cell tropism of human and avian influenza A viruses. **A)** Representative flow cytometry gating strategy of differentiated primary human bronchial epithelial cells (BEpC; donor 2). The cells were stained for the following cell surface markers: microtubules for ciliated cells, CD271 for basal cells, and CD66c for secretory cells. Cellular doublets were excluded by size, and dead cells with a cell viability marker. **B)** The relative cell type compositions of the bronchial cells from donors 1-3 were determined using the flow cytometry panel and gating strategy shown in A). Unstained or double-positive cells were categorized as ‘undefined’. **C)** BEpC from donors 1 or 2 were infected with H1N1 A/Duck/Alberta/35/1976 or A/Hawaii/70/2019, respectively, with a multiplicity of infection (MOI) of 1, 5, or 10. At 8 hours post-infection (hpi), the cells were fixed and stained for viral nucleoprotein (anti-NP antibody), tight junctions (anti-ZO-1 antibody), and nuclei (DAPI). The infections were quantified by microscopy as the percentage of NP-positive cells relative to the total number of nuclei imaged. **D)** BEpC from donors 1-3 were infected for 8 hours with H1N1 A/Duck/Alberta/35/1976 or A/Hawaii/70/2019 (MOI of 5). The cells were prepared for flow cytometry with the panel and gating strategy shown in A). The infected cells were defined as surface HA-positive cells. The relative cell type compositions are shown as the percentage of each cell type in the total populations and within the infected populations. For each donor, the values were normalized to the average across the experimental repeats. Data from donors 1-3 were pooled. One repeat of each infection was previously published in (47). **E)** The normalized infection rates of the samples described in D), as well as of BEpC from donor 1 infected under identical conditions with H1N1 Netherlands/602/2009 or Pennsylvania/02/2021. The infection rates represent the percentage of infected ciliated or secretory cells normalized to the total infected cell population. Data from donors 1-3 were pooled and distinguished by symbol shape: donor 1 (square), donor 2 (triangle), donor 3 (circle). **F)** BEpC from donor 2 were infected at an MOI of 3 with the following influenza A viruses: H1N1 Hawaii/70/2019, H3N2 Tasmania/503/2020, H3N2 Darwin/6/2021, H11N6 Duck/England/1/1956, or H3N8 Duck/Ukraine/1/1963. At 7hpi, the cells were fixed and stained for viral nucleoproteins (anti-NP antibody), ciliated cells (microtubule dye), tight junctions (anti-ZO-1 antibody), and nuclei (DAPI). The infections were quantified as the percentage of NP-positive cells relative to all apical cells and normalized to the total infected cell population. **G)** BEpC from donor 2 were infected with Hawaii/70/2019 or Duck/Alberta/35/1976 (MOI 5). At 7hpi, cells were prepared for flow cytometry and stained for infected cells (NP-positive cells). The median fluorescence intensity (MFI) of the viral NP staining in infected cells was quantified. In B) - C) and F) - G), data represent means ± s.d. from n≥3 independent experiments, and in D) - E), data represent means ± s.d. from n = 3 independent experiments per donor. In E) and F), statistical significance was determined using two-way ANOVA with Šídák’s multiple comparisons test. In G), statistical significance was determined using an unpaired t-test. *P ≤ 0.05, **P ≤ 0.01, ***P ≤ 0.001, ****P ≤ 0.0001.

To further characterize the cellular tropism during the initial round of viral infection at a transcriptomic level, we performed single-cell RNA sequencing (scRNAseq). Bronchial epithelial cells from donors 1-3 were infected for a total of 6 hours with A/Hawaii/70/2019, A/Duck/Alberta/35/1976, or a mock control. To determine the cell subtypes present in the cultures based on their transcriptomes, we used known canonical cell cluster markers defined as a transcript with at least a 0.25-fold difference (*log*-scale) and an adjusted *p*-value < 0.05 between the cells in the tested cluster and the rest of the cells. We distinguished the following epithelial cell subtypes in the cultures: ciliated, secretory, intermediate, suprabasal, basal cells, and ionocytes (Fig. 2a; Supp. Fig. 2a- b). All cell subtypes, except suprabasal cells, which were only detected in the bronchial cultures from donor 1, were detected across all three donors (Fig. 2b). The cells were further defined as infected when they expressed transcripts from at least 4 out of the 8 viral genome segments, based on previous findings showing that over 95% of genuinely influenza A infected cells meet this threshold (28) (Fig. 2c). Using this criterion, both viruses displayed consistent infection levels across donors, with around 8% infected cells for the avian strain and 10-14% for the human strain (Fig. 2d). The infection composition revealed that the human strain infected roughly equal proportions of ciliated and secretory cells (Fig. 2e), whereas the avian strain infected a larger fraction of ciliated and a smaller fraction of secretory cells. Consistent with our previous findings, only a small percentage of the basal and suprabasal cell populations were infected by both viruses. The normalized infection rates revealed a highly significant preference for ciliated cells for the avian virus and a statistically nonsignificant trend for ciliated cells for the human virus (Fig. 2f). Thus, these results are consistent with the flow cytometry and microscopy data (Fig. 1e-f). Additionally, we found that the overall copy number of viral mRNA in the hIAV-infected samples was 10-fold higher than in the aIAV-infected samples (Fig. 2g), highlighting differences in viral RNA production efficiency.

**Figure 2.**
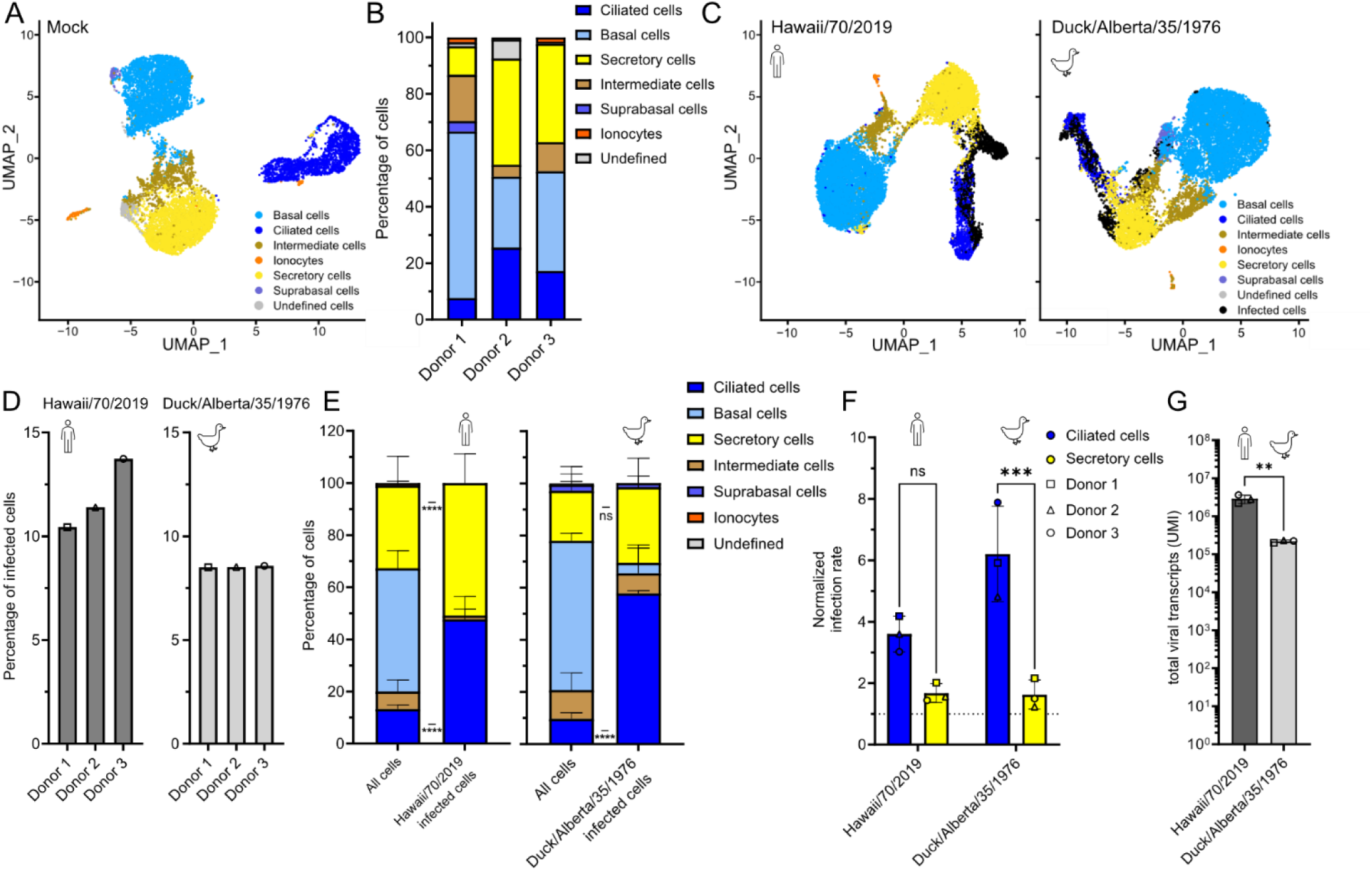
Single-cell RNA transcriptomics analysis reveals divergent bronchial epithelial cell tropism of human and avian influenza A viruses. **A)** Uniform Manifold Approximation and Projection (UMAP) visualization of single-cell RNA sequencing (scRNAseq) data from pooled mock-infected BEpC from donors 1-3. Cell subtype allocation was based on the expression of canonical subtype-specific markers, with a minimum of three markers (two for suprabasal cells) required for cluster allocation. Basal cells (light blue): TP63, KRT5, DAPL1, KRT15, ITGA6, KRT17; Secretory cells (yellow): SCGB3A1, SCGB1A1, MUC5B, MUC5AC, SPDEF, TCN1, BPIFB1, SPRR3, AGR2; Ciliated cells (dark blue): FOXJ1, CAPS, TP73, CCDC78; Ionocytes (orange): CFTR, ASCL3, FOXI1, ATP6V1C2. Suprabasal cells (violet): KRT6A and KRT15. Intermediate cells (brown): clusters expressing markers of basal and secretory cells. Undefined cells (grey): clusters not assignable to known epithelial subtype. **B)** Relative cell-type composition of BEpC from donors 1-3 of the samples described in A). **C)** UMAP visualization of the pooled scRNAseq data of BEpC from donors 1-3 infected with H1N1 Hawaii/70/2019 or Duck/Alberta/35/1976 for 6 hours with an MOI of 1. Highlighted are the allocated cell subtypes as described in A) and the infected cell populations (black). Infected cells were defined by the presence of transcripts from at least four of the eight viral genome segments. **D)** Total percentage of infected cells of the samples described in C). **E)** The relative cell type compositions of the total and the infected cell populations from the samples described in C). **F)** The normalized infection rates of the samples mentioned in C). The rates represent the percentage of infected ciliated or secretory cells normalized to the total infected cell population. **G)** Total viral transcripts detected in the samples described in C). Unique molecular identifiers (UMI). In F) and G), data from donors 1-3 were pooled and distinguished by symbol shape: donor 1 (square), donor 2 (triangle), donor 3 (circle). In A) - G), data represent n=1 experiments per donor (donor 1-3). In E) - G), data represent means ± s.d.. In E) and F), statistical significance was determined using two-way ANOVA with Šídák’s multiple comparisons test. In G), statistical significance was determined using an unpaired t-test. *P ≤ 0.05, **P ≤ 0.01, ***P ≤ 0.001, ****P ≤ 0.0001.

### The incompatibility with human ANP32 usage underlies the reduced avian influenza A virus RNA levels

In the next step, we aimed to understand the viral and host determinants underpinning the differences in viral mRNA and protein levels between hIAV and aIAV within 6 to 8 hpi in primary human airway cultures (Fig. 1g; Fig. 2g). Using reverse transcription quantitative PCR (RT-qPCR) analysis of the viral M segment levels (6 hpi), we confirmed higher viral RNA levels in the hIAV-infected over aIAV-infected samples (Fig. 3a). Using pseudobulk expression profiles of the scRNAseq data presented in Fig. 2, we performed a differential expression analysis comparing Hawaii/70/2019 infected cells over Duck/Alberta/35/1976 infected cells (6 hpi; Fig. 3b). We found that the main hits overrepresented in aIAV infected cells can be assigned to the antiviral innate immune response (Fig. 3c), with a strong presence of interferon-stimulated genes (ISGs; Fig. 3b). To test for a potential link between overrepresentation of ISGs and decreased viral RNA/protein levels in aIAV-infected cells, we inhibited the interferon-stimulated pathway and examined its effects on viral replication. We pretreated BEpC with ruxolitinib, a Janus kinase inhibitor, or control, before infecting with Hawaii/70/2019 or Duck/Alberta/35/1976. As validated by the transcript levels of IFIT2, ISG15, and MX1, we successfully inhibited the up-regulation of these ISGs at 6 hpi (Fig. 3d-f). Yet, the viral M segment levels of neither IAV were significantly affected by the presence of ruxolitinib (Fig. 3g). To test whether the presence of ruxolitinib impacts viral growth in multi-cycle replication, we harvested apically released viruses from these cultures between 1 and 48 hpi (Fig. 3h-i) and observed a significant positive effect on avian viral release (at 24hpi). These results combined highlight that while the avian virus already leads to a stronger ISG response during early BEpC infection, this does not account for the difference in viral replication at 6 hpi. However, the interferon response impacts viral growth in subsequent infection rounds.

**Figure 3.**
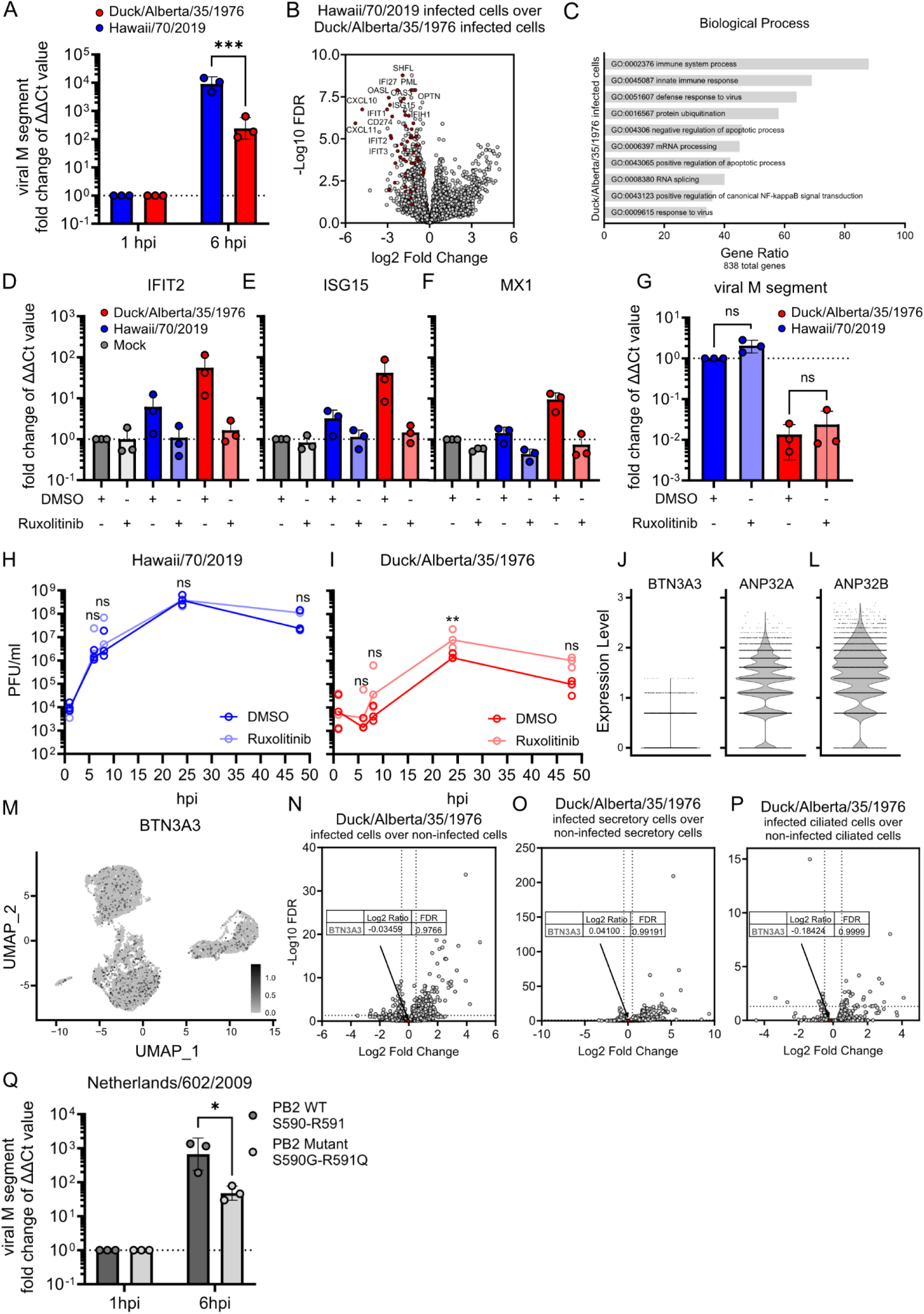
Early influenza A virus replication efficiency is dependent on ANP32 compatibility. **A)** BEpC from donors 1-3 were infected with Hawaii/70/2019 or Duck/Alberta/35/1976 (MOI 1). At 1- and 6-hours post-infection (hpi), the RNA was extracted and analyzed by RT-qPCR. The ΔCt was calculated using 18s-RNA for normalization. The enrichment of the viral M segment transcript level (ΔΔCt) is shown as a fold change relative to the 1hpi time point for each donor. **B)** and **C)** BEpC from donors 1-3 were infected with H1N1 Hawaii/70/2019 or Duck/Alberta/35/1976 for 6 hours (MOI 1) and analyzed by single-cell RNA sequencing (scRNAseq) as described in Fig. 2 C). B) The scRNAseq data were pooled to generate pseudobulk expression profiles, and a differential expression analysis was performed of Hawaii/70/2019 infected cells over Duck/Alberta/35/1976 infected cells. Genes belonging to the gene ontology GO:0045087 (innate immune response) are highlighted (dark red). C) A hypergeometric over-representation analysis (ORA) of the differentially expressed genes (DEG) identified in B). The plot shows the top 10 enriched biological processes, ranked with the number of genes associated with each process overrepresented in the Duck/Alberta/35/1976 infected cell population. The threshold for DEG was set at *P ≤ 0.01. **D)** - **G)** BEpC from donor 2 were pretreated with DMSO (control) or ruxolitinib for 86 hours prior to infection. The cells were infected with mock, Hawaii/70/2019, or Duck/Alberta/35/1976 (MOI 1). At 6 hpi, the RNA was extracted and analyzed by RT-qPCR. The ΔCt was calculated using 18s-RNA for normalization. The enrichment (ΔΔCt) of the transcript levels of IFIT2, ISG15, MX1, and viral M segment are presented as a fold change relative to D) – F) mock DMSO control or G) Hawaii/70/2019 DMSO control. **H)** – **I)** BEpC from donor 2 were pretreated and infected as described for D). At the indicated time points post-infection, the apical viral release was harvested and plaqued in MDCK cells. **J) – L)** Mock-infected BEpC from donors 1-3 were prepared for scRNAseq, as described in Fig 2 A). Plots depicting single cell transform (SCTransform) -normalized expression of J) BTN3A3, K) ANP32A, and L) ANP32B within all cells from the bronchial mock-infected cells (donor 1-3 pooled). **M)** UMAP visualization of the BTN3A3 SCTransform-normalized expression within the bronchial mock-infected cells (donor 1-3 pooled). **N)** – **P)** Differential expression analysis was performed of Duck/Alberta/36/1976 N) infected cells over non-infected cells, O) infected secretory cells over non-infected secretory cells, and P) infected ciliated cells over non-infected ciliated cells. BTN3A3 is highlighted (dark red). **Q)** BEpC from donor 2 were infected with Netherlands/602/2009 PB2 S590-R591 or with Netherlands/602/2009 PB2 S590G-R591Q (MOI of 0.6). At 1 and 6 hpi, the RNA was extracted and analyzed by RT-qPCR. The ΔCt was calculated using 18s-RNA for normalization. The enrichment of the viral M segment transcript level (ΔΔCt) is shown as a fold change relative to the 1hpi time point for each donor. In A) - C) and J) – P) data represent n=1 experiments per donor (donor 1-3; means ± s.d.). In D) – G) and Q) data represent means ± s.d. from n=3 independent experiments. In H) – I), data represent medians from n=3 independent experiments. In A), G) – I), and Q), statistical significance was determined using two-way ANOVA with Šídák’s multiple comparisons test. *P ≤ 0.05, **P ≤ 0.01, ***P ≤ 0.001.

We next aimed to assess the role of known human host factors that can either promote (hANP32A and hANP32B) (12) or inhibit (BTN3A3) (17) IAV replication, with differential effects on human versus avian IAV. While the human IAV codes for the PB2 SR polymorphism at the amino acid positions 590 and 591 and is therefore able to efficiently use the hANP32 proteins to replicate its genome (29,30), it also codes for NP 313V and thus can escape BTN3A3 restriction (17) (Supp. Fig. 3a). On the other hand, the avian virus does not carry substitutions known to enable hANP32 usage or BTN3A3 escape. Transcript analysis of the mock-infected cultures confirmed the expression of the three genes (Fig. 3j-l). The ANP32 transcripts showed high expression across the cell population, whereas BTN3A3 showed a comparably low expression profile. As BTN3A3 expression was restricted to a subset of cells (Fig. 3m), we hypothesized that BTN3A3 expression would be overrepresented in non-infected cells if it was responsible for the observed difference of aIAV RNA levels. However, no preference for infection of BTN3A3-negative cells was detected, even upon restriction of the analysis to a specific cell subtype population (ciliated or secretory cells) (Fig. 3n-p), suggesting that BTN3A3 is not the root of the difference in early viral RNA levels between hIAV and aIAV in the primary human airway cultures. Next, to determine whether the viral polymerase adaptation to hANP32 usage underlies the differences in viral replication efficiency, we generated a recombinant H1N1 virus based on the Netherlands/602/2009 strain in which the PB2 segment was mutated to encode the hANP32-maladapted R590G-S591Q sequence (Supp. Fig. 3a). Using RT-qPCR analysis, we found the loss of the PB2 SR polymorphism significantly impaired replication efficiency at 6hpi (Fig. 3q), highlighting the importance of the host factor ANP32 for early viral replication.

### Influenza A virus tropism in human epithelial cells is linked to the haemagglutinin protein and the apical distribution of sialic acid linkages

To determine the basis of the cellular tropism of A/Hawaii/70/2019 and A/Duck/Alberta/35/1976, we first examined viral determinants and generated recombinant 7+1 viruses. These viruses carry the internal segments (PB2, PB1, PA, NP, M, and NS) of the laboratory-adapted H1N1 strain A/WSN/1933 and express either the envelope protein HA or neuraminidase (NA) of either Hawaii/70/2019 or Duck/Alberta/35/1976 (Fig. 4a; Supp. Fig. 4a). At 7hpi, the parental strain A/WSN/1933 showed no preference for infecting either ciliated or apical non-ciliated cells. Likewise, exchanging the neuraminidase did not alter tropism. However, substitution of HA for the avian variant was sufficient to induce a clear and significant preference for ciliated over non-ciliated bronchial epithelial cells. These infection patterns directly correlate with the tropism seen in the flow cytometry (Fig. 1e), microscopy (Fig. 1f), and scRNAseq (Fig. 2f) data. Given that HA mediates viral attachment and entry via sialic acid residues on the cell surface, we examined the distribution of α2,3- or α2,6-linked sialic acids on the whole surface of BEpC (pan-membrane staining). Using the flow cytometry panel for the different cell types together with lectins for either α2,3- or α2,6-linked sialic acids (Fig. 4b-g; Supp. Fig. 4b), we found that the percentages of α2,3- or α2,6-linked sialic acid-positive cells did not significantly differ between ciliated and secretory cells (Fig. 4b and e; BEpC from donors 1-3). Likewise, the median fluorescent intensity (MFI) of these stainings, when considering all ciliated and secretory cells, did not differ significantly between the populations (Fig. 4c and f). However, when analyzing the signal intensity specifically on sialic acid-positive cells (Fig. 4d and g), we found that the α2,3-linked sialic acid signal was consistently higher on ciliated versus secretory cells (Fig. 4d), giving a first indication that ciliated cells display a higher surface density of α2,3-linked sialic acids than α2,3-linked sialic acid-positive secretory cells.

**Figure 4.**
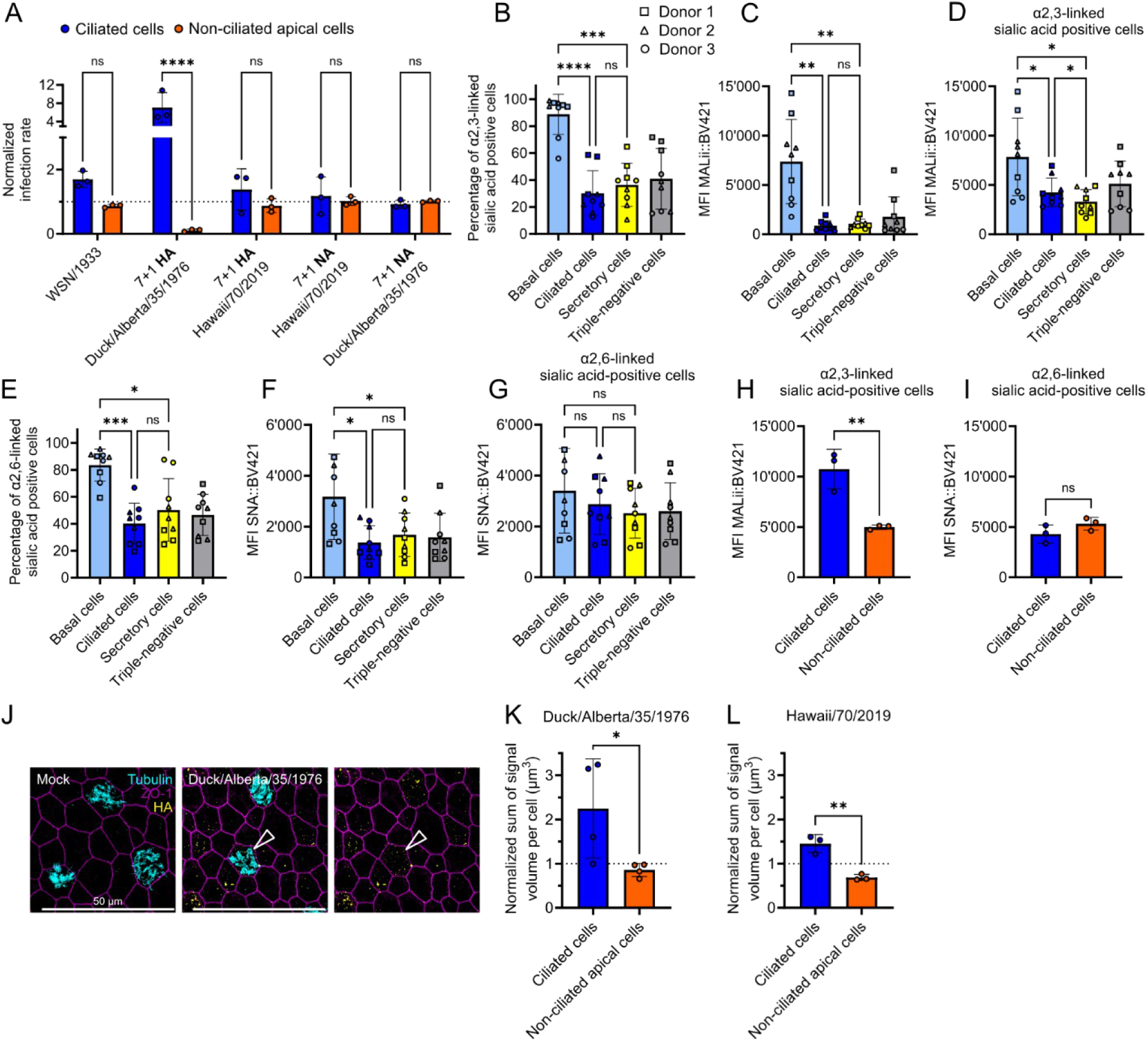
The bronchial epithelial cell tropism of influenza A virus is mediated by haemagglutinin and the apical distribution of sialic acid linkages. **A)** BEpC from donor 2 were infected with control H1N1 WSN/1933 virus or recombinant 7+1 viruses containing 7 genes from the WSN/1933 strain and either the HA- or NA-encoding segment of Hawaii/70/2019, or of Duck/Alberta/35/1976. An MOI as calculated by titres determined on MDCK cells of 0.5-1 was used for the recombinant strains, and an MOI of 5 was used for the control WSN/1933 strain to achieve comparable infection rates. At 7 hours post-infection (hpi), the cells were fixed and stained for viral nucleoproteins (anti-NP antibody), ciliated cells (microtubule dye), tight junctions (anti-ZO-1 antibody), and nuclei (DAPI). The infections were quantified as the percentage of NP-positive cells relative to all apical cells and normalized to the total infected cell populations. **B)** – **G)** BEpC from donor 1-3 were trypsinized and stained for cell surface markers identifying ciliated cells (microtubules), basal cells (CD271), secretory cells (CD66c), α2,3-linked sialic acids (Maackia amurensis II lectins; MALII), and α2,6-linked sialic acids (Sambucus nigra lectins; SNA). The cells were analyzed by flow cytometry, cellular doublets were excluded by size, and dead cells with a cell viability marker. One repeat was previously published in (36). B) and E) show the percentage of α2,3-linked and α2,6-linked sialic acid-positive cells, respectively. C) and F) show the median fluorescent intensity (MFI) of the BV421 staining of α2,3-linked and α2,6-linked sialic acids, respectively. D) and G) show the MFI of the BV421 staining of α2,3-linked and α2,6-linked sialic acids, respectively, of pre-gated sialic acid-positive cells. The data of donors 1-3 were pooled and distinguished by symbol shape: donor 1 (square), donor 2 (triangle), donor 3 (circle). **H)** – **I)** BEpC from donor 2 were stained prior to trypsinization for apical H) α2,3-linked sialic acids (MALII) and I) α2,6-linked sialic acid (SNA), as well as ciliated cells (microtubule dye). The cells were analyzed by flow cytometry, cellular doublets were excluded by size, and dead cells with a cell viability marker. Shown is the MFI of the apical sialic acid staining on ciliated versus non-ciliated cells, pre-gated for sialic acid-positive cells. **J)** - **L)** BEpC from donor 2 were precooled on ice, and 6.4*10^9 viral copies of K) Duck/Alberta/35/1976 or L) Hawaii/70/2019 were added apically on ice. 1 hour post addition, the cells were fixed and stained for apically bound viral particles (anti-HA antibody), ciliated cells (microtubule dye), tight junctions (anti-ZO-1 antibody), and nuclei (DAPI). Quantification of apically bound viral signal. The summed viral signal volume on ciliated and non-ciliated apical cells was normalized to the viral signal present on all apical cells. J) Representative microscopy images of the mock-treated and Duck/Alberta/35/1976-treated sample are shown. The white arrows highlight the viral HA staining on a cell, shown with and without the cilia staining. In A), and H) - L), data represent means ± s.d. from n≥3 independent experiments. In B) – G) data represent means ± s.d. from n = 3 independent experiments per donor (donor 1-3). Statistical significance was determined in A) using two-way ANOVA with Šídák’s multiple comparisons test, in B) – G) using RM one-way ANOVA with Tukey’s multiple comparisons test, and in H) – I) and K) – L) using an unpaired t-test. *P ≤ 0.05, **P ≤ 0.01, ***P ≤ 0.001, ****P ≤ 0.0001.

As the viruses are restricted to the apical surface of the cells for the initial binding and entry, we confined further staining to the apical surface only. Microscopy analysis confirmed the presence of both α2,3- and α2,6-linked sialic acids on the apical surface of ciliated and non-ciliated cells (Supp. Fig. 4c). Furthermore, flow cytometry enabled quantitative assessment of their levels (Fig. 4h-i; Supp. Fig. 4d). While the MFI was comparable between α2,6-linked sialic acid-positive ciliated and non-ciliated cells, the α2,3-linked sialic acids were detected at significantly higher levels on ciliated cells versus non-ciliated cells. These data highlight that, although levels of α2,3-linked sialic acid on the whole cell surface barely differed between ciliated and secretory cells, we identified a clear disparity on the apical cell surface. To further delineate the origin of the cellular tropism, we next assessed viral binding to the airway cultures. The viruses were allowed to attach to the apical cell surface of the bronchial human airway cells on ice, after which the viral particles were stained for microscopy analysis (Fig. 4j-l). Notably, the sum of the viral signal per cell was significantly higher on ciliated than on non-ciliated apical cells for both viruses (Fig. 4k-l). For the avian virus, this is in line with our finding of higher α2,3-linked sialic acid levels on ciliated cells. For the human virus, the binding pattern does not correlate with the α2,6-linked sialic acid levels, suggesting that glycan characteristics beyond linkage and/or additional attachment receptors affect virus binding.

### Receptor preference of avian H1N1 for sialyl Lewis X correlates with glycan abundance on ciliated cells

To determine if glycan structures besides the terminal linkage of the sialic acid residue play a role in the binding efficiency of the viruses, we conducted glycan array analysis for Hawaii/70/2019 and Duck/Alberta/35/1976 (Fig. 5a; Supp. Fig. 5a-d) (31,32). As expected, the avian strain showed a clear preference for α2,3-linked sialic acids (Fig. 5b), while the human strain preferred binding to α2,6-linked sialic acids (Fig. 5e). Beyond the linkages, we were able to reveal a preference of the avian IAV for longer and more branched glycans (Fig. 5b; Supp. Fig. 5a) and a preference for sialylation of specific glycan arms over others (Fig. 5b; Supp Fig. 5b). Specifically, the avian virus prefers sialylation on the mannosyl-glycoprotein N-acetylglucosaminyl transferase 4 (MGAT4) over the MGAT5 antenna. Lastly, aIAV displayed a clear preference for sialyl Lewis X (sLe^x^) epitopes over non-fucosylated α2,3-linked sialic acid residues (Fig 5b; Supp. Fig. 5c). Via microscopy and anti-sLe^x^ antibodies, we could confirm the presence of sLe^x^ on the apical surface of ciliated and non-ciliated cells (Fig. 5c). Further, via flow cytometry, we quantified the signal of apical sLe^x^ and identified 3-times higher levels on ciliated versus non-ciliated cells (Fig. 5d; Supp. Fig. 5e), highlighting that the preference of the avian H1N1 strain for sLe^x^ correlates with the glycan presence on the apical epithelium surface. The human IAV, on the other hand, showed a preference for two N-acetyllactosamine (LacNAc) repeats on the sialylated branch over one or three repeats (Fig. 5f-g), when both branches of a bi-antennary N-glycan are sialylated. In addition, the human virus also bound bi-antennary glycans where one branch had 3 LacNAc repeats and the other branch only one LacNAc repeat (Fig. 5g; Supp. Fig. 5d). From these glycan array results combined with the viral binding data, we speculate that ciliated cells display more of the bi-antennary glycans that are preferred by the human virus, while overall α2,6-linked sialic acid levels are comparable between ciliated and secretory cells.

**Figure 5.**
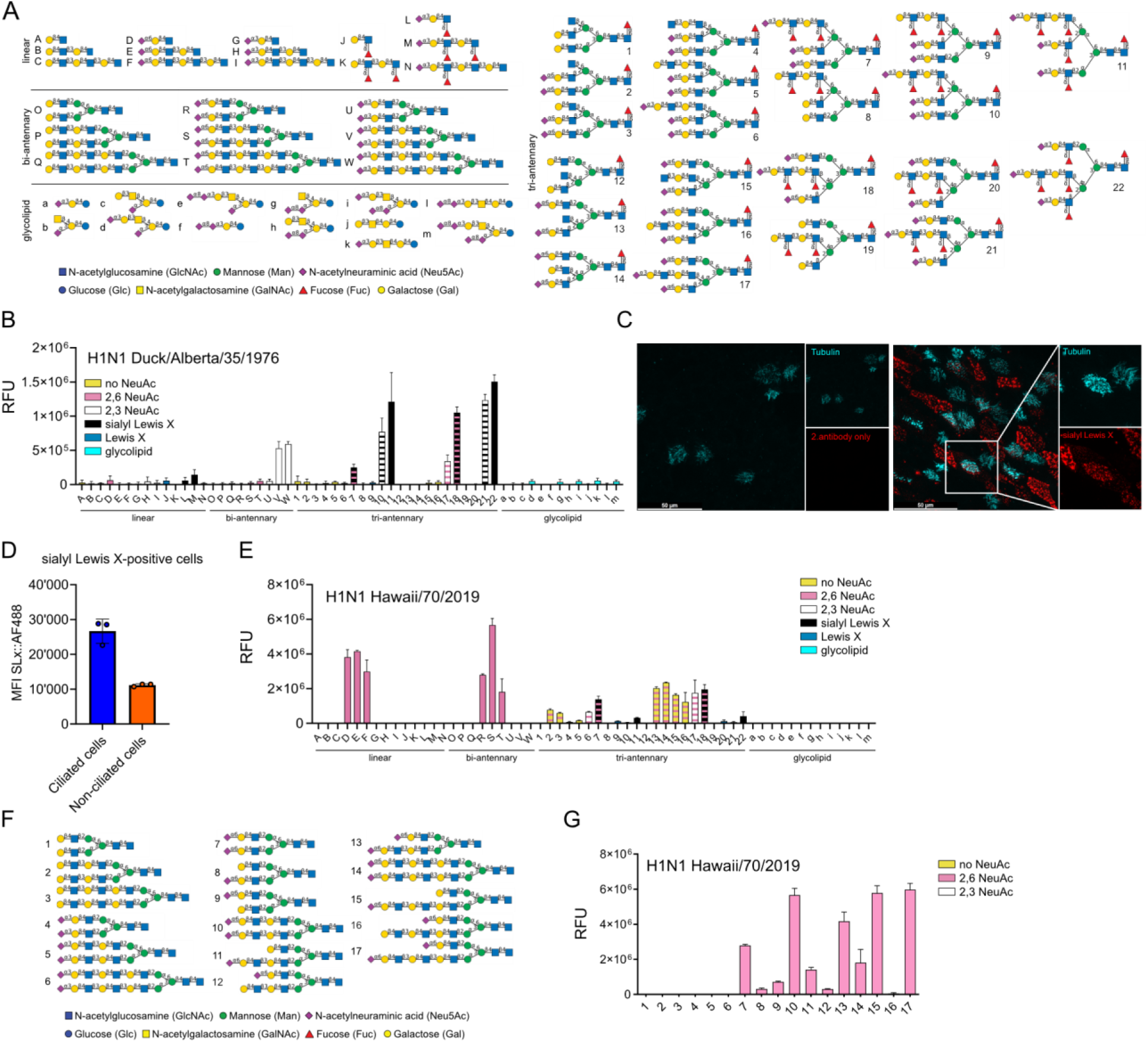
avian H1N1 preference for sialyl Lewis X correlates with glycan abundance on ciliated cells. **A)** Synthetic glycans on the microarray N°1 with linear (A-N), bi-antennary (O-W), tri-antennary structures (1–22), and glycolipids (a-m). **B)** Microarray N°1 binding of H1N1 A/Duck/Alberta/35/1976. The microarray N°1 tested for glycans with no sialylation (yellow), α2,6-linked NeuAc (pink), α2,3-linked NeuAc (white), Lewis X (blue), sialyl Lewis X (dark blue), and glycolipids (cyan). Striped bars indicate glycans terminating in different epitopes on different arms. **C)** Representative microscopy images of apical sialyl Lewis X staining on BEpC from donor 2. The cells were stained for ciliated cells (microtubule dye) and surface sialyl Lewis X. **D)** BEpC from donor 2 were stained prior to trypsinization for ciliated cells (microtubule dye), as well as for apical sialyl Lewis X (sLx). The cells were analyzed by flow cytometry, cellular doublets were excluded by size, and dead cells with a cell viability marker. Shown is the MFI of the apical sialyl Lewis X staining on ciliated versus non-ciliated cells, pre-gated for sLx-positive cells. **E)** Microarray N°1 binding of A/Hawaii/70/2019. **F)** Synthetic glycans on the microarray N°2. **G)** Microarray N°2 binding of H1N1 A/Hawaii/70/2019. The microarray N°2 tested for glycans with no sialylation (yellow), α2,6-linked NeuAc (pink), α2,3-linked NeuAc (white). In B), E), and G), mean relative fluorescence units (RFU) ± s.d. from n = 4 technical repeats are shown. In D), data represent means ± s.d. from n=3 replicates.

### Early viral replication levels of human H1N1 are cell subtype specific

Given the significant preference for ciliated cells in binding but not overall infection rate for the human strain, we aimed to test for differences in transcription/replication levels between ciliated and secretory cells. First, we employed our scRNAseq analysis (Fig. 2) and quantified viral mRNA copies in both cell types. Indeed, viral mRNA levels were significantly higher in infected secretory versus infected ciliated cells for Hawaii/70/2019 but not for Duck/Alberta/35/1976 (Fig. 6a). In parallel, we also measured significantly higher viral NP protein levels in non-ciliated versus ciliated cells for the Hawaii/70/2019 infected but not for the Duck/Alberta/35/1976 infected cells (Fig. 6b). Based on these findings, we investigated whether known host factors contribute to the virus- and cell type-specific differences in replication efficiency. Using pseudobulk expression profiles of the scRNAseq data, we performed differential expression analysis comparing mock secretory over ciliated cells (Fig. 6c). Gene ontology (GO) analysis of the transcripts enriched in each cell type revealed that ciliated cells were primarily associated with cilium-related processes (Fig. 6d), whereas secretory cells showed an enrichment for pathways related to transcription including positive regulation of RNA polymerase II (GO:0045944) (Fig. 6e). Given that previous work has established that IAV transcription depends on RNA polymerase II activity (33–36), these findings suggest that viral transcription may be more efficient in secretory cells due to higher active RNA polymerase II activity. To assess whether cell-type-specific, infection-induced transcriptional responses, such as the type I interferon response, contribute to the difference in viral replication, we compared the induction of infection-induced genes relative to the mock controls (Fig. 6f). As the top infection-induced transcripts exhibited consistently higher fold changes in secretory cells than ciliated cells, we concluded that the cell-type-specific infection responses, including the ISG expression levels, cannot explain the increase in RNA production observed in secretory cells. Lastly, the comparison of ANP32 expression revealed that secretory cells expressed approximately 25% higher levels of ANP32B and 17% higher levels of ANP32A (Fig. 6g-h). Using the Netherlands/602/2009 viral pair, we found that the ability to utilize human ANP32 proteins not only influences overall replication efficiency (Fig. 3q), but also accounts for the higher viral protein levels observed in secretory compared to ciliated cells (Fig. 6i; Supp. Fig. 6a-b). It therefore highlights the importance of the host factor ANP32 in early viral replication and cellular tropism.

**Figure 6.**
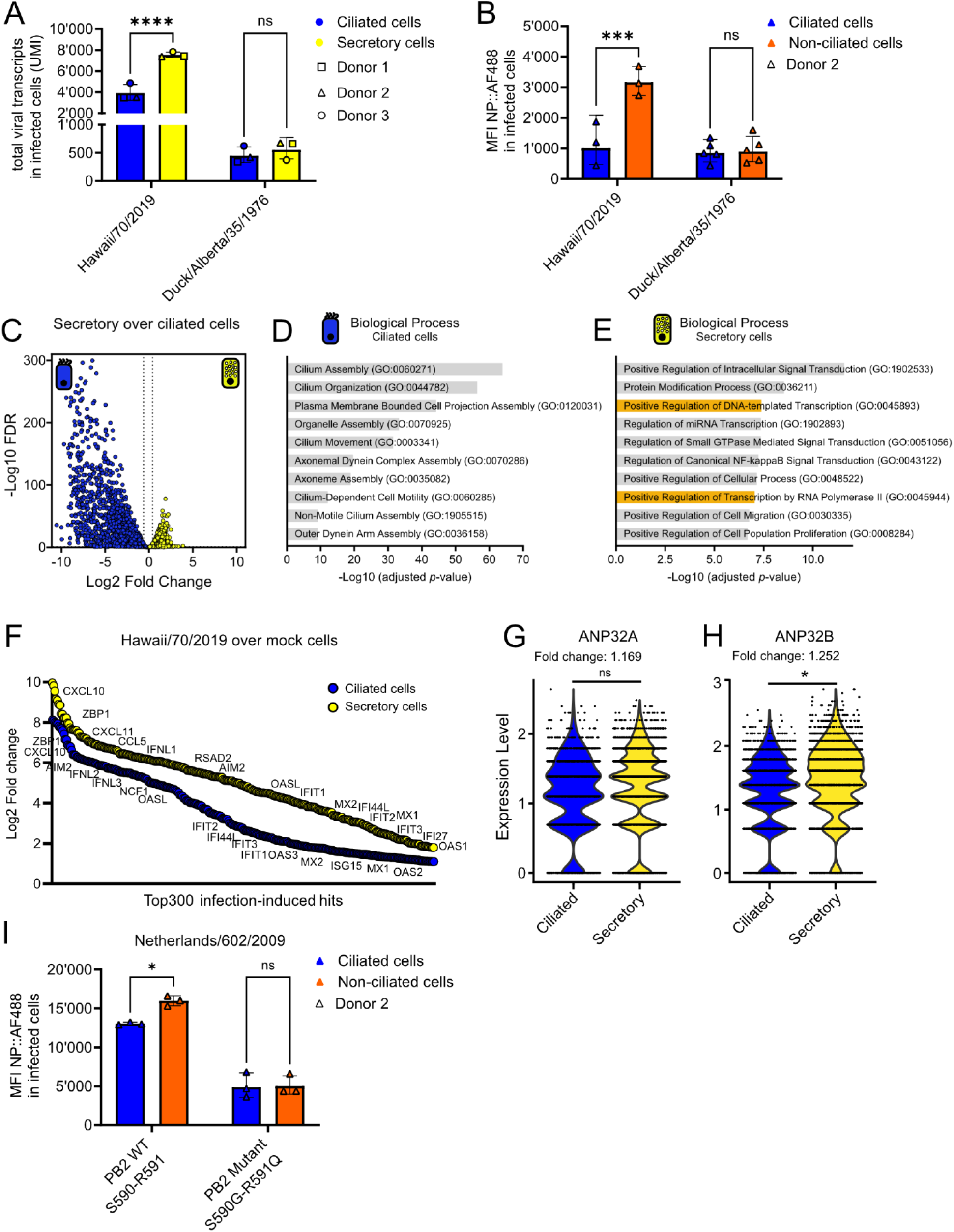
Early viral RNA and protein levels of human H1N1 are cell subtype specific. **A)** BEpC from donors 1-3 were infected with H1N1 Hawaii/70/2019 or Duck/Alberta/35/1976 for 6 hours (MOI1) and analyzed by single-cell RNA sequencing (scRNAseq), as described in Fig. 2 C). Total unique molecular identifiers (UMI) detected in infected secretory and ciliated cells. Data from donors 1-3 were pooled and distinguished by symbol shape: donor 1 (square), donor 2 (triangle), donor 3 (circle). **B)** BEpC from donor 2 were infected with Hawaii/70/2019 or Duck/Alberta/35/1976 (MOI 5; 7hpi) and prepared for flow cytometry as described in Fig 1 G). The cells were stained for infected cells (NP-positive cells) and ciliated cells (microtubule staining). The median fluorescence intensity (MFI) of the viral NP staining in infected ciliated and non-ciliated cells was quantified. **C)** The scRNAseq mock data was pooled to generate pseudobulk expression profiles, and a differential expression analysis of mock secretory cells over mock ciliated cells was performed. **D)** – **E)** A gene ontology analysis of the differentially expressed genes (DEG) identified in C). The plot shows the top 10 enriched biological processes, ranked by the adjusted *p*-value that were overrepresented in D) the ciliated cell and E) the secretory cell population. The threshold for DEG was set at *P ≤0.05, and an absolute log2 fold change of 0.5. **F)** The scRNAseq data was pooled to generate pseudobulk expression profiles, and a differential expression analysis of Hawaii/70/2019 infected samples over mock samples was performed for ciliated and secretory cell. **G)** – **H)** Single cell transform (SCTransform) -normalized expression of G) ANP32A and H) ANP32B within each subtype of the scRNAseq data set from bronchial mock-infected cells. **I)** BEpC from donor 2 were infected with Netherlands/602/2009 PB2 S590-R591 or with Netherlands/602/2009 PB2 S590G-R591Q (MOI of 0.45). At 8hpi, cells were prepared for flow cytometry and stained for infected cells (NP-positive cells) and ciliated cells (microtubule staining). The median fluorescence intensity (MFI) of the viral NP staining in infected ciliated and non-ciliated cells was quantified. In A) and C) – H), data represent n =1 experiments per donor (donor 1-3). The scRNAseq mock data was pooled to generate pseudobulk expression profiles, and a differential expression analysis of mock secretory cells over mock ciliated cells was performed to calculate the fold change and the statistical significance. In B) and I), data represent n ≥3 independent experiments. In A), B), and I), data represent means ± s.d. and statistical significance was determined using two-way ANOVA with Šídák’s multiple comparisons test. *P ≤ 0.05, **P ≤ 0.01, ***P ≤ 0.001.

In sum, our results show that human and avian IAV infect ciliated and secretory cells in the human airway epithelium. Avian IAV displays a strong preference for ciliated cells, which is driven by the higher abundance of sLe^x^ on ciliated apical cell surfaces and thus stronger viral binding. Combined with comparable replication efficiencies in both cell types, avian IAV displays an overall preference for ciliated cells. In contrast, the tropism of human IAV is driven by both differential viral binding and differential RNA levels across the two cell types. While binding is stronger on ciliated cells, possibly via higher abundance of bi-antennary glycans with two LacNac repeats, viral RNA and protein production are more efficient in secretory cells due to differential ANP32A/B expression. This leads to an overall dual tropism for both cell types for human IAV.

## DISCUSSION

A zoonotic spillover from the avian IAV reservoir into the human host is initiated by a productive infection in the airway epithelium, where early host-pathogen interactions determine the outcome. Our study demonstrates that such infections exhibit distinct tropism patterns compared to those of seasonal human influenza viruses. Importantly, our findings highlight that the cellular tropism is not dictated by individual pro- or antiviral factors in isolation but instead arises from the interplay of multiple viral and host determinants acting across different stages of the viral life cycle.

Using a differentiated human airway epithelial model, we provide a comprehensive system to resolve the heterogeneity of IAV infection at the cellular level, enabling direct comparison between human and avian IAV infections as well as stratification by epithelial cell subtypes. Through this approach, we redefine IAV tropism in the human airway and show that at least two key steps in the viral life cycle (viral binding and viral RNA production) shape the outcome with stage-specific differences that are not apparent when only assessing the overall infection.

For viral binding, we show that the overall glycan structure influences it, not solely sialic acid linkages. While it has been reported before that certain avian IAV strains preferentially bind sialyl Lewis X (sLe^x^) (H5, H6, H7, and H9) (31,37,38), we now show that this preference may be more widespread, demonstrating preferential binding of an avian H1N1 virus to sLe^x^. Moreover, we uncover that sLe^x^ is enriched on ciliated apical cell surfaces, the primary target of avian viruses in our model. On the other hand, human-adapted H3N2 strains are known to have evolved to bind to extended (≥3 LacNAc repeats) branched glycans (39–43). Human H1N1 strains have been shown to preferentially bind sialylated glycans with two LacNAc repeats (31,42,43), which is the preference that we have identified for our human H1N1 strain. By combining pan-membrane and apical-surface-specific analyses, we uncover pronounced differences in glycan distribution not only across epithelial cell subtypes but also between staining approaches. Whereas previous studies relied on non-quantitative immunofluorescence (18,22,23) or focused exclusively on pan-membrane staining (24), our quantitative approach reveals distinct apical versus pan-membrane patterns of α2,3-linked sialic acids. This provides a more refined, cell subtype-specific view of the epithelial glycome than previously reported. These findings emphasize that both spatial and cell subtype-specific glycan presentation can influence viral tropism.

For viral replication, we find that the host factors ANP32A/B represent a major limitation for avian IAV within the initial replication round in the primary airway cultures. This highlights the importance of polymerase compatibility as a barrier during early zoonotic infection and is in line with previous reports that studied ANP32A/B function in more conventional cell lines (12,29,30). Additionally, the preferential replication of seasonal human viruses in secretory cells suggests that ciliated cells may represent a suboptimal environment for viral replication and that differential expression of ANP32A/B contributes to this observation. Taken together, our data indicate that viral entry is not the primary limiting step, but that efficient replication represents a key bottleneck in this model.

Lastly, our findings also provide an important context for previous work. Roach *et al.* (2024) (24) reported that pandemic 2009 strains produce significantly higher levels of viral transcripts in bronchial secretory cells compared to ciliated cells at 6 hours post-infection. We extend this analysis by identifying the role of ANP32 in this process, and by comparing avian IAV infections within the same system and reveal that the increased viral RNA levels in secretory cells are specific for human IAV. We therefore underscore fundamental differences in host adaptation.

It is important to note that while the differentiated primary cell cultures provide a powerful system to study the heterogeneity and complexity of the human airway, they also show limitations. In particular, the current difficulty in genetically manipulating these cultures restricts mechanistic investigations. This is especially apparent when aiming to study cell subtype-specific aspects. For example, it remains challenging to dissect the correlation between human viral binding preferences and specific N-glycan structures directly in such complex systems.

In summary, our study provides a comprehensive view of influenza A virus tropism that integrates both extracellular attachment and intracellular replication determinants. We show that cellular susceptibility is shaped not only by the glycan landscape but also by cell type-specific replication capacity, supporting a multifactorial model of early IAV infection and zoonotic transmission. Together, these findings represent a conceptual advance in our understanding of influenza virus host adaptation: by combining a physiologically relevant model with a stage-resolved analysis of infection, this work moves beyond receptor-centric paradigms and identifies intracellular compatibility as a critical determinant of zoonotic success.

## METHODS

### Cell lines

Human embryonic kidney cells (HEK) 293T (CRL-3216™) and Madin-Darby canine kidney (MDCK) cells (CCL-34™) were purchased from the American Type Culture Collection (ATCC). The cells were cultured at 37°C, 5% CO2, and >80% relative humidity in Dulbecco’s modified Eagle’s medium (DMEM; Gibco, #41966-052) supplemented with 10% heat-inactivated fetal calf serum (FCS; Gibco, #A5256701) and 100 U/mL of Penicillin-Streptomycin (Gibco, #15140-122).

### Primary human epithelial cultures

Primary human bronchial epithelial cells (BEpC) were purchased from Epithelix (#EP51AB). Cells from three independent donors were used: BEpC_AB079 (donor 1; male, 62 years old, Hispanic, non-smoker), BEpC_AB051 (donor 2; male, 53 years old, Caucasian, non-smoker), BEpC_AB0839 (donor 3; male, 59 years old, Caucasian, non-smoker). Cells were grown in airway epithelium basal growth medium (Promocell, #C-21260) supplemented with the airway epithelial cell growth medium supplement pack (Promocell, #C-39160) and 10 µM Y-27632 (ROCK inhibitor; Tocris, #1254). For cellular differentiation, the transwell plates with 12 mm (Corning, #CLS3460) or 6.5 mm (Corning, #CLS3470) filter inserts were coated with collagen. The human collagen type 1 (Sigma-Aldrich, #C7774) stock in 0.5 M acetic acid (Merck, #100063.1000) was diluted to 0.15 mg/ml in PBS before coating the filters. The primary cells were seeded onto the filter inserts in a 1:1 mixture of airway epithelium basal growth medium and DMEM, which was supplemented with airway epithelial cell growth medium supplement pack (Gray’s medium). The cells were grown until confluency was reached. For the cellular differentiation at the air-liquid interface (ALI), Gray’s medium was removed from the apical side of the filter inserts, and Gray’s medium in the basal compartment was supplemented with 150 ng/mL retinoic acid (Sigma-Aldrich, #R2625). The cells were cultured at ALI for a minimum of 28 days prior to use. Medium was refreshed in the basal compartment every 2-3 days, and epithelial integrity was monitored weekly by determining the transepithelial electrical resistance (TEER) using an ERS-2 meter (Millicell, #MERS00002).

### Viruses

The IAV strains H1N1 A/Duck/Alberta/35/1976, H3N8 A/Duck/Ukraine/1/1963, and H11N6 A/Duck/England/1/1956 were grown in 10-day-old embryonated chicken eggs. H1N1 A/Hawaii/70/2019 was grown in primary human bronchial epithelial cells. H1N1 A/Pennsylvania/02/2021, H3N2 A/Tasmania/503/2020, H3N2 A/Darwin/6/2021, H1N1 A/Netherlands/602/2009, and H1N1 A/WSN/1933 were grown on MDCK cells. Virus stocks were titrated by plaque assay on MDCK cells. Virus sequences were verified by next-generation sequencing.

### Infection of primary human epithelial cells

BEpC were washed three times (for a total of 15 minutes at 37°C) with Dulbecco’s phosphate-buffered saline (PBS; Gibco, #14190144) to remove the mucus from the apical side of the inserts. Cells were infected with the indicated viral strain at the indicated multiplicity of infection (MOI) in infection PBS (PBSi; PBS supplemented with 0.3% Bovine Serum Albumin solution (BSA; Sigma-Aldrich, #A1595), 1 mM Ca2+/Mg2+, and 100 U/mL of Penicillin-Streptomycin (Gibco, #15140-122)). After incubation for 1 hour at 37°C, the inoculum was removed, and the apical side of the insert was washed once with PBS.

### Microscopy analysis of viral binding on primary human bronchial cultures

BEpC were washed three times (for a total of 15 minutes at 37°C) with PBS (Gibco, #14190144) to remove the mucus from the apical side of the inserts. Cells were stained for cilia, using the SiR-tubulin kit (1:1,000; Cytoskeleton/Spirochrome, #CY-SC002) for 1 hour at 37°C in Gray’s medium on the apical side of the filter inserts. The cells were washed once with PBS and precooled on ice for 15 minutes. 6.4*10^9^ viral copies/insert were added apically to the cells on ice. At 1 hour post viral addition, the unbound virus was washed off and the cells were fixed with 4% paraformaldehyde solution (PFA; Electron Microscopy Sciences, #15710) in PBS and blocked for unspecific staining with blocking buffer N°1 (BB1; PBS supplemented with 3% BSA (Sigma-Aldrich, #A7906)) for 30 minutes at RT. The viral particles were stained using an anti-HA 1.12 antibody (1:100) (44) for 1 hour at RT. The unbound antibodies were washed off 3-times with BB1 before permeabilizing the cells for 15 minutes with the permeabilization buffer N°1 (PB1; PBS supplemented with 50mM ammonium chloride (Sigma-Aldrich, #254134), 0.1% saponin (Sigma-Aldrich, #47036), and 2% BSA (Sigma-Aldrich, #A7906)). Next, cells were stained with the primary antibody against tight junctions anti-ZO-1 (1:50; Thermo Fisher Scientific, #61-7300) for 1 hour at RT. The unbound antibodies were washed off 3-times with PB1, before the cells were stained with the secondary antibodies: Alexa Fluor™ 546 donkey anti-rabbit IgG highly cross-adsorbed secondary antibody (1:1,000; Invitrogen, #A10040) and Alexa Fluor™ 488 goat anti-human IgG Fc recombinant secondary antibody (1:500; Invitrogen, #A55747). The nuclei were stained with 4′,6-Diamidino-2-phénylindole dihydrochloride (DAPI) (1:1,000; Roche, #10236276001). The cell-containing membranes were mounted on a microscope slide by using ProLong™ gold antifade Mountant (Thermo Fisher Scientific, #P36930). Cells were imaged using the DMi8 microscope (Leica, Germany) and processed using the THUNDER large volume computational clearing algorithm (Leica). Maximum projection images of z-stacks were generated using the LAS X software (Leica). The images were analysed using Imaris 10.2 (Oxford Instruments) to quantify the viral signal on the surface of ciliated and non-ciliated cells.

### Microscopy analysis of infected primary human bronchial cultures

BEpC were infected with the respective viruses and MOIs as indicated previously. 1 hour before fixation, the cells were stained for cilia, using the SiR-tubulin kit (1:1,000; Cytoskeleton/Spirochrome, #CY-SC002) for 1 hour at 37°C in Gray’s medium on the apical side of the filter inserts. The cells were washed once with PBS and then fixed with 4% PFA (Electron Microscopy Sciences, #15710) in PBS. The samples were permeabilized with permeabilization buffer N°1 (PB1; PBS supplemented with 50mM ammonium chloride (Sigma-Aldrich, #254134), 0.1% saponin (Sigma-Aldrich, #47036), and 2% BSA (Sigma-Aldrich, #A7906)) for 30 minutes at RT. Next, cells were stained with the primary antibodies: anti-NP (1:5; ATCC, #HB-65) and anti-ZO-1 (1:50; Thermo Fisher Scientific, #61-7300) for 1 hour at RT. When indicated, the cells were additionally stained with an anti-HA 1.12 antibody (1:100) (44). The unbound antibodies were washed off 3-times with PB1, before cells were stained with the secondary antibodies: Alexa Fluor™ 546 donkey anti-rabbit IgG highly cross-adsorbed secondary antibody (1:1,000; Invitrogen, #A10040), and Alexa Fluor™ 488 goat anti-mouse IgG highly cross-adsorbed secondary antibody (1:1,000; Invitrogen, #A-11029). The nuclei were stained with DAPI (1:1,000; Roche, #10236276001). The cell-containing membranes were mounted on a microscope slide by using ProLong™ gold antifade Mountant (Thermo Fisher Scientific, #P36930). Cells were imaged using the DMi8 microscope (Leica, Germany) and processed using the THUNDER large volume computational clearing algorithm (Leica). Maximum projection images of z-stacks were generated using the LAS X software (Leica). A minimum of 1,000 total/apical cells were counted per sample. The total percentages of infected cells were quantified for all cells containing an apical surface in the selected images (or, if mentioned, for all nuclei present in the selected images). All apical cells negative for the cilia staining were grouped as ‘non-ciliated cell with an apical surface’.

### Flow cytometry analysis of infected primary human bronchial cultures (cell surface staining)

The flow cytometry protocol was developed based on the protocol from Bonser *at al.* (2021) and previously described in Karakus *et al.* (2024) (27,45). In brief, BEpC were infected with respective viruses and MOIs as indicated previously. At 1 hour prior to trypsinization, cells were stained for cilia, using the SiR-tubulin kit (1:1,000; Cytoskeleton / Spirochrome, #CY-SC002) for 1 hour at 37°C in Gray’s medium on the apical side of the filter inserts. Cells were washed once with PBS and trypsinized by adding 0.25% Trypsin-EDTA (Gibco, #25200-056). Cells were resuspended in warm DMEM/F12 (Gibco, #11330-032) supplemented with 5% FCS (Gibco, #A5256701). The resuspended cells were washed once with PBS (805 x*g* for 5 minutes at 4°C) and dead cells were stained using the LIVE/DEAD fixable near-IR dead cell marker (1:1,000; Thermo Fisher Scientific, #L10119) in PBS for 30 min at 4 °C. The marker was washed away once in PBS. Subsequently, cells were fixed in 4% PFA (Electron Microscopy Sciences, #15710) in PBS and blocked with blocking buffer N°2 (BB2; PBS supplemented with 5% normal goat serum (NGS; Gibco, #16210-064)) for ∼15 minutes at room temperature (RT). Cells were stained with the following cell surface markers: for secretory cells using BV510 mouse anti-human CD66c antibodies (1:100; BD Biosciences, #742684), for basal cells using PE/Cyanine7 anti-human CD271 (NGFR) antibodies (1:50; BioLegend, #345110), for α2,3-linked sialic acids using Maackia amurensis II lectin (MALII; 1:50) (Vector laboratories, #B-1265-1), for α2,6-linked sialic acids using Sambucus nigra lectin (SNA; 1:425) (Vector laboratories, #B-1305) plus Brilliant Violet 421™ Streptavidin (1:200; BioLegend, #405226), and infected cells using an anti-HA 1.12 antibody (1:100) (44) plus PE conjugated anti-human IgG Fc secondary antibody (1:100; Invitrogen, #12-4998-82). Cells were washed and resuspended in FACS buffer (FB, PBS supplemented with 2% BSA (Sigma-Aldrich, #A7906) and 1 mM EDTA (Thermo Fisher Scientific, #AM9260G)). Cells were analysed either with the BD FACSVerse flow cytometer (BD Biosciences) and the BD FACSuite™ application or with the BD FACSymphony™ A1 Cell Analyzer (BD Biosciences) and the BD FACSDiva™software. A minimum of 10,000 total events were acquired per sample. Data analysis was done using the FlowJo v.10 software (BD Life Sciences).

### Flow cytometry analysis of infected primary human bronchial cultures (intracellular staining)

BEpC were infected with respective viruses and MOIs as indicated previously. At 1 hour before trypsinization, the cells were stained for cilia, using the SiR-tubulin kit (1:1,000; Cytoskeleton / Spirochrome, #CY-SC002) for 1 hour at 37°C in Gray’s medium on the apical side of the filter inserts. Cells were washed once with PBS and trypsinized by adding 0.25% Trypsin-EDTA (Gibco, #25200-056). The cells were resuspended in warm DMEM/F12 (Gibco, #11330-032) supplemented with 5% FCS (Gibco, #A5256701). The resuspended cells were washed once with PBS (805 x*g* for 5 minutes at 4°C), before staining dead cells using the LIVE/DEAD fixable near-IR dead cell stain (1:1,000; Thermo Fisher Scientific, #L10119) in PBS for 30 min at 4 °C. The marker was washed away once in PBS. Subsequently, cells were fixed in 4% PFA (Electron Microscopy Sciences, #15710) in PBS and permeabilized with permeabilization buffer N°2 (PB2; PBS supplemented with 2% BSA (Sigma-Aldrich, #A7906), 1 mM EDTA (Thermo Fisher Scientific, #AM9260G), and 0.1% (w/v) saponin (Sigma-Aldrich, #47036)). The infected cells were stained with anti-NP antibodies (1:5; ATCC, #HB-65) for 1 hour at RT in BB2. The unbound antibodies were washed off twice, and the cells were stained with the Alexa Fluor™ 488 goat anti-mouse IgG highly cross-adsorbed secondary antibody (1:500; Invitrogen, #A-11029) for 1 hour at RT in BB2. The cells were washed and resuspended in FB. The cells were analysed either with the BD FACSVerse flow cytometer (BD Biosciences) and the BD FACSuite software or with the BD FACSymphony™ A1 Cell Analyzer (BD Biosciences) and the BD FACSDiva™ software. A minimum of 10,000 total events were acquired per sample. Data analysis was done using the FlowJo v.10 software (BD Life Sciences).

### Single-cell RNA sequencing processing of infected primary human bronchial cultures

BEpC donor 1-3 were infected with either H1N1 A/Hawaii/70/2019, H1N1 A/Duck/Alberta/35/1976, or mock for a total of 6 hours at an MOI of 1, as indicated above. The cells were then washed once and trypsinized by adding 0.25% Trypsin-EDTA (Gibco, #25200-056) for a maximum of 10 minutes at 37°C and resuspended in warm DMEM/F12 (Gibco, #11330-032) supplemented with 5% FCS (Gibco, #A5256701). Cells were centrifuged at 400 x*g* for 5 minutes at 4°C and washed once in cold PBS supplemented with 0.04% BSA (Sigma-Aldrich, #A7906). Subsequently, cells were resuspended in cold 0.04% BSA in PBS and filtered through a 40 μm Flowmi cell strainer (Merck, #BAH136800040) to ensure single-cell suspension. Using the Countess II automated cell counter (ThermoFisher), a final live cell concentration was measured at 600-1,200 cells/μL. Next, using the Chromium Controller (10X Genomics) and following the ‘Chromium Next GEM Single Cell 3′ reagent kits v.3.1 (Dual Index)’ protocol (10X Genomics), the 3′ gene expression library was constructed. For sequencing, the libraries were processed on an Illumina NovaSeq X Plus flow cell. The sequencing parameters were set according to 10X Genomics recommendations. The Functional Genomics Centre Zurich performed sequencing and subsequent computational data analysis.

### Single-cell RNA sequencing SUSHI pipeline

Single-cell RNA sequencing data analysis was performed using the SUSHI framework (46), which encompassed the following steps. A chimeric reference genome was constructed using the R package ezRun (v3.19.1) (47) by supplementing the human genome assembly (GRCh38.p13, GENCODE Release 42) (48) with the nucleotide sequences of the two influenza A virus strains used in this study: H1N1 A/Hawaii/70/2019 and H1N1 A/Duck/Alberta/35/1976. Sample demultiplexing, alignment of reads against this chimeric reference, and generation of feature–barcode count matrices were performed using 10x Genomics Cell Ranger v7.2.0 (49), with pre-mRNA counting enabled (--include-introns=true) to capture unspliced transcripts. Downstream analysis was performed on the resulting count matrices using the R package Seurat v5.1.0 (50) under R version 4.4.0 and Bioconductor 3.19. Ambient RNA contamination was estimated and removed using DecontX (celda v1.20.0) (51) and SoupX (v1.6.2) (52). Doublets were identified and removed using scDblFinder (53). Low-quality cells were filtered using a median-absolute-deviation (MAD) approach implemented in the R package scater (54): cells deviating by more than 3 MADs below the median library size or number of detected genes, or more than 3 MADs above the median mitochondrial read fraction, were excluded. Cells with ribosomal protein gene content exceeding 70% were additionally removed. Genes expressed in fewer than 0.01% of cells were filtered out before downstream analysis. Normalisation and variance stabilisation of count data were performed using SCTransform v2 (sctransform v0.4.1) (55), computed on the top 3,000 highly variable genes. Dimensionality reduction was performed by principal component analysis (PCA), and the top 20 principal components were retained for downstream steps. Cell clusters were identified using the Louvain algorithm at a resolution of 0.6. Cluster marker genes were determined by Wilcoxon rank-sum test, retaining genes with an average log₂ fold-change ≥ 0.4, Bonferroni-adjusted p-value < 0.01, expressed in at least 10% of cells in the tested cluster, and with a minimum difference in detection rate of 10% between the tested cluster and all other cells. Cell subtype allocation was based on the expression of the canonical subtype-specific markers, with a minimum of three canonical markers (two for suprabasal cells) required for cluster allocation. Basal cells: TP63, KRT5, DAPL1, KRT15, ITGA6, KRT17; Secretory cells: SCGB3A1, SCGB1A1, MUC5B, MUC5AC, SPDEF, TCN1, BPIFB1, SPRR3, AGR2; Ciliated cells: FOXJ1, CAPS, TP73, CCDC78; Ionocytes: CFTR, ASCL3, FOXI1, ATP6V1C2. Suprabasal cells: KRT6A and KRT15. Intermediate cells: clusters expressing markers of basal and secretory cells. Undefined cells: clusters not assignable to known epithelial subtype. Donor-level batch correction and integration were performed separately for each experimental condition (mock, Hawaii/70/2019-infected, Duck/Alberta/35/1976-infected) using Harmony (56) as implemented within the SUSHI ScSeuratCombine workflow, combining the three donors within each condition. Infected cells were defined as those expressing a minimum of four viral gene segments, where a segment was considered expressed if its UMI count exceeded 1.5 times the mean expression of that segment across all cells in the same sample. Per-cell viral transcript abundance was computed as log_10_(1 + summed viral UMIs) and summarised per cell type as the mean of these per-cell values within each group.

### Pseudobulk dataset

Pseudobulk and differential expression analysis was performed using the SUSHI framework (46). Pseudobulk count matrices were generated by summing raw UMI counts per cell type per sample from the individually processed, cell type-labelled per-sample Seurat objects, treating each sample as a biological replicate. Differential expression between conditions was tested using DESeq2 (57) as implemented in the SUSHI twoGroups workflow, under R version 4.4.0 and Bioconductor 3.19. Gene ontology (GO) enrichment was performed using the clusterProfiler Bioconductor R package (58) against the GO Biological Process, Molecular Function, and Cellular Component ontologies. Over-representation analysis (ORA) was applied to genes reaching a DESeq2 p-value ≤ 0.05 and absolute log₂ fold-change > 0.5; significant GO terms were defined at FDR ≤ 0.05.

### Generation of recombinant viruses

to generate recombinant IAVs, the viral genes coding for HA and NA from Hawaii/70/2019 and Duck/Alberta/35/1976 were cloned into ambisense expression plasmids (pDZ) as previously described (45). In brief, viral RNA was extracted from the virus stock using the QIAamp Viral RNA mini kit (Qiagen, #52904) according to the manufacturer’s protocol. The viral segments were amplified by PCR with the Q5® High-Fidelity PCR kit (NEB, #E0555S) using universal primers (Supp. Table 1) and cloned into the expression vector pDZ using the restriction enzyme SapI (NEB, #R0569S). XL10-Gold ultracompetent cells were used for transformation (Agilent, #200315). The expression plasmids of the WSN/1933 genes were a gift from Benjamin G. Hale (Universität Zürich, Switzerland). The reverse genetics system for Netherlands/602/2009 was previously described in Medina *et al*., 2013 (59). For the mutation of the PB2 plasmid of the Netherlands/602/2009 strain, the segment was amplified with the targeted mutation using designed primer pairs (Supp. Table 1) and cloned into the pDZ plasmid via In-Fusion cloning (Takara Bio, #638948). Stellar™ competent cells were used for transformation (Thermo Fisher Scientific, #NC9905817)

The recombinant viruses were rescued as previously described (60). In brief, HEK-293T cells were transfected with 0.5 ug of the following 8 plasmids: PB2, PB1, PA, HA, NP, NA, M, and NS-encoding segments of the respective viruses using FuGENE® HD transfection reagent (Promega, #E2312) in OPTI-iMEM® (Gibco, #31985062). At 24 hours post-transfection, MDCK cells (or MDCK expressing chicken ANP32 (61)) were co-seeded with 1 ug/ml TPCK-trypsin (Sigma-Aldrich, #T1426) for additional 48 hrs. The rescued viruses were plaque-purified and further propagated in MDCK cells (or MDCK expressing chicken ANP32). Viral sequences were verified by next-generation sequencing, and viral titres were determined by plaque assay on MDCK cells.

### Apical surface glycan staining for immunofluorescence

Mucus was removed by washing with PBS. For the sialidase treatment, cells were preincubated at 37°C for 1 hour with 400 mU/ml neuraminidase from *Vibrio cholerae* (Sigma-Aldrich, #N6514-1UN). For cilia staining, cells were incubated with the SiR-tubulin kit (1:1,000; Cytoskeleton / Spirochrome, #CY-SC002) for 1 hour at 37°C in Gray’s medium on the apical side of the filter inserts. Cells were fixed with 4% PFA (Electron Microscopy Sciences, #15710) in PBS and blocked for unspecific staining with BB1. The apical surface alpha-2,3-linked and alpha-2,6-linked sialic acid linkages were stained using biotinylated Maackia amurensis II lectin (MALII; 1:50) (Vector laboratories, #B-1265-1) or Sambucus nigra lectin (SNA; 1:425) (Vector laboratories, #B-1305) in BB1 for 1 hour at RT. The apical surface sialyl Lewis X (sLe^x^; 1:100) were stained (BD Biosciences, #551344 (clone CSLEX1)) in BB1 for 1 hour at RT.

The lectins/antibodies were washed off 3-times with BB1 before permeabilizing the cells for 15 minutes with PB1. For tight junction staining, cells were incubated with an anti-ZO-1 antibody (1:50; Thermo Fisher Scientific, #61-7300). For cilia staining in combination with sialic acid staining, an anti-beta IV tubulin antibody (1:400; Abcam, #ab11315) was used in PB1 for 1 hour at RT. The antibodies were washed away 3-times with PB1. Lastly, cells were incubated with secondary proteins/antibodies for 1 hour at RT: streptavidin Alexa Fluor™ 647 conjugate (1:500; Invitrogen, #S21374), Alexa Fluor™ 546 donkey anti-rabbit IgG highly cross-adsorbed secondary antibody (1:1,000; Invitrogen, #A10040), Alexa Fluor™ 488 goat anti-mouse IgG highly cross-adsorbed secondary antibody, (1:1,000; Invitrogen, #A-11029), and Alexa Fluor™ 488 goat anti-mouse IgM cross-adsorbed secondary antibody (1:500; Thermo Fisher Scientific, #A-21042). Nuclei were stained with DAPI (1:1,000; Roche, #10236276001). The cell-containing membranes were mounted on a microscope slide by using ProLong™ gold antifade mountant (Thermo Fisher Scientific, #P36930). The cells were imaged using the DMi8 microscope (Leica, Germany) and processed using the THUNDER large volume computational clearing algorithm (Leica). Maximum projection images of z-stacks were generated using the LAS X software (Leica).

### Apical surface glycan staining for flow cytometry

Mucus was removed from the bronchial donor 2 cells. The cells were stained for cilia, using the SiR-tubulin kit (1:1,000; Cytoskeleton / Spirochrome, #CY-SC002) for 1 hour at 37°C in Gray’s medium on the apical side of the filter inserts. The cells were precooled on ice and blocked for unspecific binding on the apical surface with BB2 on ice. The apical surface of the live cells were stained as described for the immunofluorescent analysis: for alpha-2,3-linked sialic acids using MALII (1:50; Vector laboratories, #B-1265) and alpha-2,6-linked sialic acid using SNA (1:425; Vector laboratories, #B-1305), or for sialyl Lewis X (1:100; BD Biosciences, #551344 (clone CSLEX1)) in BB2 for 1 hour on ice. The unbound lectins/antibodies were washed off 3-times with BB2. Next, cells were stained with Brilliant Violet 421™ streptavidin (1:200; BioLegend, #405226) or with Alexa Fluor™ 488 goat anti-mouse IgM cross-adsorbed secondary antibody (1:500; Thermo Fisher Scientific, #A-21042) for 30-60 minutes on ice. The unbound proteins were washed off 3-times with BB2 and once with PBS on the apical and the basolateral side. Cells were trypsinized by adding 0.25% Trypsin-EDTA (Gibco, #25200-056) and resuspended in warm DMEM/F12 (Gibco, #11330-032) supplemented with 5% FCS (Gibco, #A5256701). Cells were then washed once with PBS (805 x*g* for 5 minutes at 4°C) before staining the dead cells using the LIVE/DEAD fixable near-IR dead cell stain (1:1,000; Thermo Fisher Scientific, #L10119) in PBS for 30 min at 4 °C. The marker was washed away once in PBS. The cells were acquired with the BD FACSymphony™ A1 Cell Analyzer (BD Biosciences), the data was collected with the FACSDiva™ software. A minimum of 10,000 total events were acquired per sample. Data analysis was done using the FlowJo v.10 software (BD Life Sciences).

### RNA extraction and reverse transcription quantitative PCR (RT-qPCR)

RNA extraction was conducted using the RNeasy Kit (Qiagen, #74181) and the QIAshredder (Qiagen, #79656) according to the manufacturer’s instructions. Reverse transcription was done using the SuperScript™ IV reverse transcriptase (Thermo Fisher Scientific, #18090050), following the manufacturer’s protocol using random primers (Promega AG, #C1181), dNTP mix (Invitrogen, #18427088), and RNasin® Ribonuclease Inhibitor (Promega AG, #N2615). The qPCR was done using PowerTrack™ SYBR Green master mix (Thermo Fisher Scientific, #A46109) using the appropriate primers (Supp. Table 1). Data was acquired using a 7300 Real-Time PCR System with a 7300 System SDS Software (Applied Biosystems). The relative gene expression was calculated with the ΔΔCt method using 18s-rRNA for normalization (62). Statistical analysis was conducted on linear ΔCt values.

### Ruxolitinib treatment

BEpC from donor 2 were pretreated basally for 86 hours prior to infection with either 5 μM ruxolitinib (Santa Cruz, #sc-364729) or 5μM Dimethyl sulfoxide (DMSO; Sigma-Aldrich, #41640-100ML) as a control.

### Viral growth curve

BEpC were infected with respective viruses and MOIs as described above. At indicated time points post infection, 60 µL of PBSi were added apically to the cells for 5 min at 37°C and then collected and stored at −80°C. Virus titers were determined by plaque assay on MDCK cells.

### Glycan array of H1N1 viruses

Viral stocks were inactivated using 0.03% PFA (Electron Microscopy Sciences, #15710) for 4 days at 4°C. For hemagglutinin (HA) titer determination, viral stocks were serially diluted 2-fold in PBS and incubated with 0.5% chicken red blood cells. An HA titer of 1:128 – 1:512 was used. Consequently, the viral isolates were diluted in PBS-T (PBS+0.1% Tween-20) and applied to subarrays in the presence of oseltamivir (200 nM) in a humidified chamber for 1 hour. Next, the microarray slide was rinsed with PBS-T, PBS, and deionized water (twice) and dried by centrifugation. The slide was incubated for 1 hour in the presence of the CR6261 stem-specific antibody (100 µL, 5 µg/mL^−1^ in PBS-T) (32) and washed as described above. A secondary goat anti-human Alexa Fluor-647 antibody (100 µL, 2 µg/mL^−1^ in PBS-T) (Thermo Fisher Scientific, #A-21445) was applied, and the resulting slide was incubated for 1 hour in a humidified chamber and then washed by the standard procedure. Afterwards, the slides were rinsed successively with PBS-T, PBS, and deionized water. Slides were dried by centrifugation after the washing steps and scanned immediately using an Innopsys Innoscan 710 microarray scanner. Various gains and PMT values were used to ensure the signals were in the linear range and to avoid saturation of the signals. Images were analyzed with Mapix software (version 8.1.0 Innopsys) and processed with an Excel macro (https://github.com/enthalpyliu/carbohydrate-microarray-processing). The average fluorescence intensity and standard deviation were determined for each compound after removal of the highest and lowest intensities from the spot replicates to give n=4.

## ACKNOWLEDGEMENTS

We thank the Functional Genomics Center Zurich for their support with the single-cell transcriptomics approach, sequencing, and data analysis. We thank Benjamin G. Hale (Universität Zürich, Switzerland) for the WSN/1933 expressing plasmids and the chicken ANP32 expressing MDCK cells (61). This study was funded by the Swiss National Science Foundation (grant number 310030_204166 to SSt) and the UZH Candoc Grant (grant number FK-25-039 to JvK).

## Notes

### Competing Interest Statement

The authors have declared no competing interest.

