## Supplementary figures for "Opposing cell type preferences for binding and replication shape influenza A virus infection in human airways"

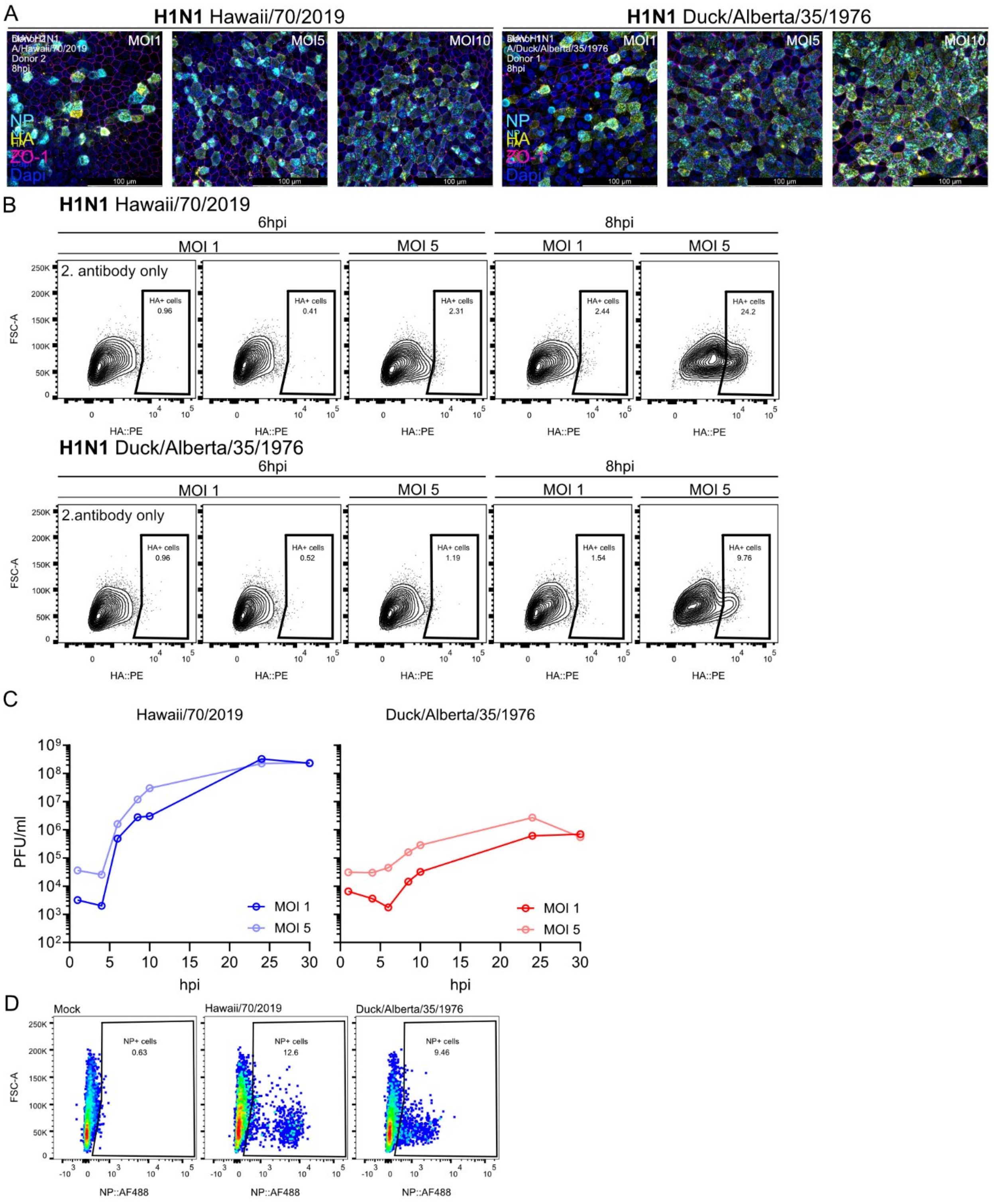


**Supp. figure 1. Flow cytometry and microscopy analysis reveal divergent bronchial epithelial cell tropism of human and avian influenza A viruses.**

**A)** Representative microscopy images of the samples described in Fig 1 C). BEpC from donors 1 or 2 were infected with H1N1 A/Duck/Alberta/35/1976 or A/Hawaii/70/2019, respectively, with a multiplicity of infection (MOI) of 1, 5, or 10. At 8 hours post-infection (hpi), the cells were fixed and stained for viral nucleoprotein (anti-NP antibody), tight junctions (anti-ZO-1 antibody), and nuclei (DAPI). **B)** BEpC from donor 3 were infected with H1N1 Duck/Alberta/35/1976 or Hawaii/70/2019 at an MOI of 1 or 5 for 6 to 8 hours. Cells were prepared for flow cytometry.

Cellular doublets were excluded by size, and dead cells using a cell viability marker. The percentage of infected cells was measured by gating for cells with surface hemagglutinin (HA) expression. Secondary antibody only staining was used as a negative control. **C**) BEpC from donor 3 were infected with H1N1 A/Duck/Alberta/35/1976 or A/Hawaii/70/2019, with an MOI of 1 or 5. At the indicated time points post-infection, the apical viral release was harvested and plaqued in MDCK cells. **D)** Representative dot plots of the gating strategy of NP-positive cells for Fig 1 G). BEpC from donor 2 were infected with Hawaii/70/2019 or Duck/Alberta/35/1976 (MOI 5). At 7hpi, cells were prepared for flow cytometry and stained for infected cells (NP-positive cells).

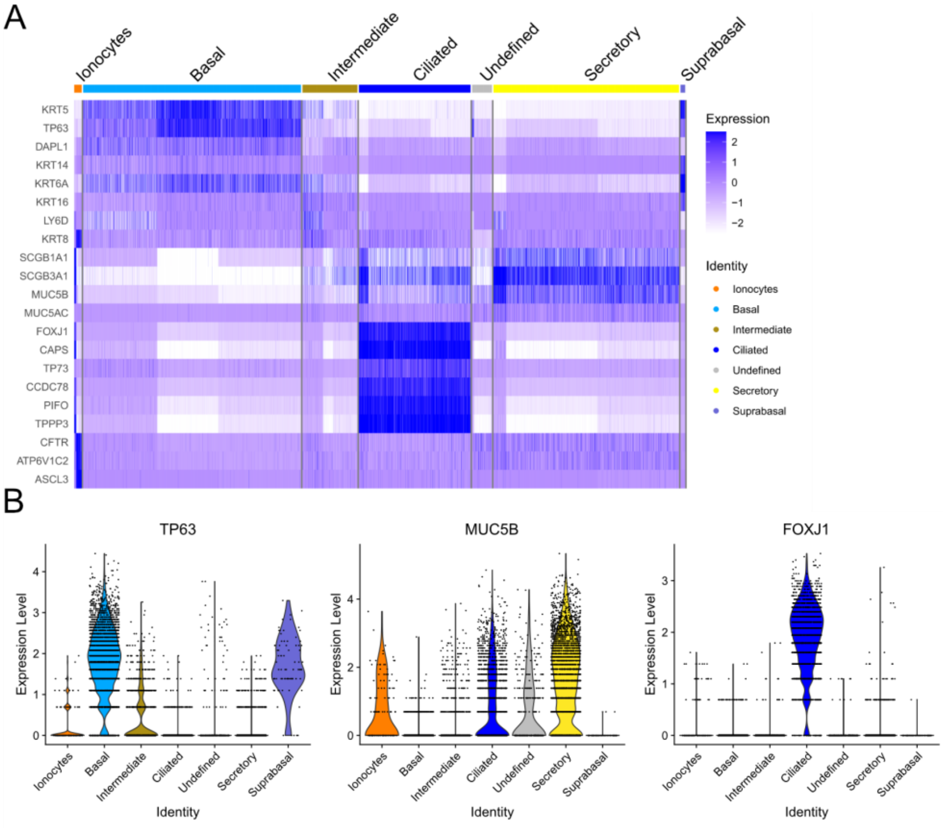


**Supp. figure 2. Defining single-cell sequencing cell cluster identity via cell cluster markers.**

**A)** Heatmap depicting single cell transform (SCTransform) -normalized expression of a set of canonical subtype-specific markers in the single-cell RNA sequencing (scRNAseq) data set from bronchial donors 1-3 mock-infected, described in Fig 2 A). Cell subtype allocation was based on the expression of canonical subtype-specific markers, with a minimum of three markers (two for suprabasal cells) required for cluster allocation. Basal cells (light blue): TP63, KRT5, DAPL1, KRT15, ITGA6, KRT17; Secretory cells (yellow): SCGB3A1, SCGB1A1, MUC5B, MUC5AC, SPDEF, TCN1, BPIFB1, SPRR3, AGR2; Ciliated cells (dark blue): FOXJ1, CAPS, TP73, CCDC78; Ionocytes (orange): CFTR, ASCL3, FOXI1, ATP6V1C2. Suprabasal cells (violet): KRT6A and KRT15. Intermediate cells (brown): clusters expressing some markers of basal and secretory cells. Undefined cells (grey): clusters not assignable to known epithelial subtype. **B)** Violin plots depicting SCTransform-normalized expression of TP63, MUC5B, and FOXJ1 within each subtype of the scRNAseq data set from bronchial mock-infected cells (donor 1-3 pooled).

A

| PB2 | 590 | 591 | 627 | 701 |
| --- | --- | --- | --- | --- |
| Netherlands/602/2009 | S | R | E | D |
| Hawaii/70/2019 | S | R | E | D |
| Duck/Alberta/35/1976 | G | Q | E | D |

| NP | 52 | 313 |
| --- | --- | --- |
| Netherlands/602/2009 | Y | V |
| Hawaii/70/2019 | Y | V |
| Duck/Alberta/35/1976 | Y | F |

**Supp. figure 3. Early influenza A virus transcription and replication efficiency is independent of interferon-stimulated gene expression.**

**A)** Amino acid sequence alignment of PB2 and NP highlighted positions (PB2 590, 591, 627, and NP 52, 313) across viruses (Netherlands/602/2009, Hawaii/70/2019, Duck/Alberta/35/1976).

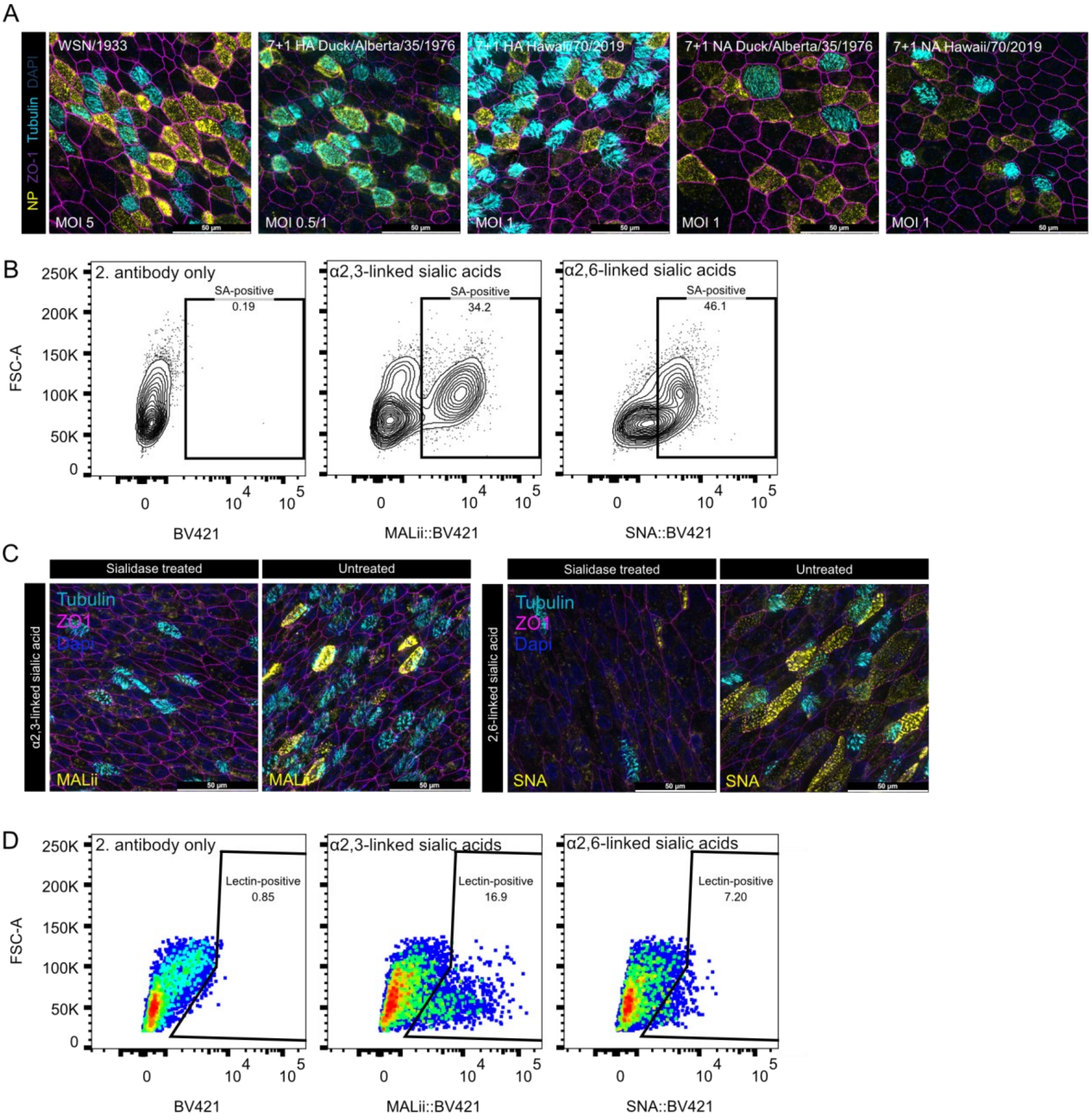


**Supp. figure 4. Influenza A virus tropism in human bronchial epithelial cells is mediated by haemagglutinin and the apical distribution of sialic acid linkages.**

**A)** Representative microscopy images of the samples described in Fig 4 A). BEpC from donor 2 were infected with control H1N1 WSN/1933 virus or recombinant 7+1 viruses containing 7 genes from the WSN/33 strain and either the HA- or NA-encoding segment of Hawaii/70/2019, or of Duck/Alberta/35/1976. A multiplicity of infection (MOI) of 0.5-1 was used for the recombinant strains, and an MOI of 5 was used for the control WSN/1933 strain to achieve comparable infection rates. 7 hours post-infection (hpi), the cells were fixed and stained for viral nucleoproteins (anti-NP antibody), ciliated cells (microtubule dye), tight junctions (anti-ZO-1 antibody), and nuclei (DAPI). **B)** Representative dot plots of gating strategy to determine sialic acid positive cells for Fig 4 B) - G). BEpC from donor 2 were trypsinized and stained for cell surface markers identifying ciliated cells (microtubules), basal cells (CD271), secretory cells (CD66c), α2,3-linked sialic acids (Maackia amurensis II lectins), and α2,6-linked sialic acids (Sambucus nigra lectins). The cells were analyzed by flow cytometry, cellular doublets were excluded by size, and dead cells with a cell viability marker. **C)** Representative microscopy images of apical sialic acid stainings on BEpC from donor 2 (untreated or sialidasetreated). The cells were fixed and stained for surface α2,3-linked sialic acids (Maackia amurensis II lectins) or α2,6-linked sialic acid (Sambucus nigra lectins), as well as for tight junctions (anti-ZO-1 antibody), ciliated cells (anti-beta IV tubulin antibody), and nuclei (DAPI). **D)** Representative dot plots of gating strategy to determine apically sialic acid positive cells for Fig 4 H) – I). BEpC from donor 2 were stained prior to trypsinization for apical α2,3-linked sialic acids (Maackia amurensis II lectins) and α2,6-linked sialic acid (Sambucus nigra lectins), as well as ciliated cells (microtubule dye). The cells were analyzed by flow cytometry, cellular doublets were excluded by size, and dead cells with a cell viability marker.

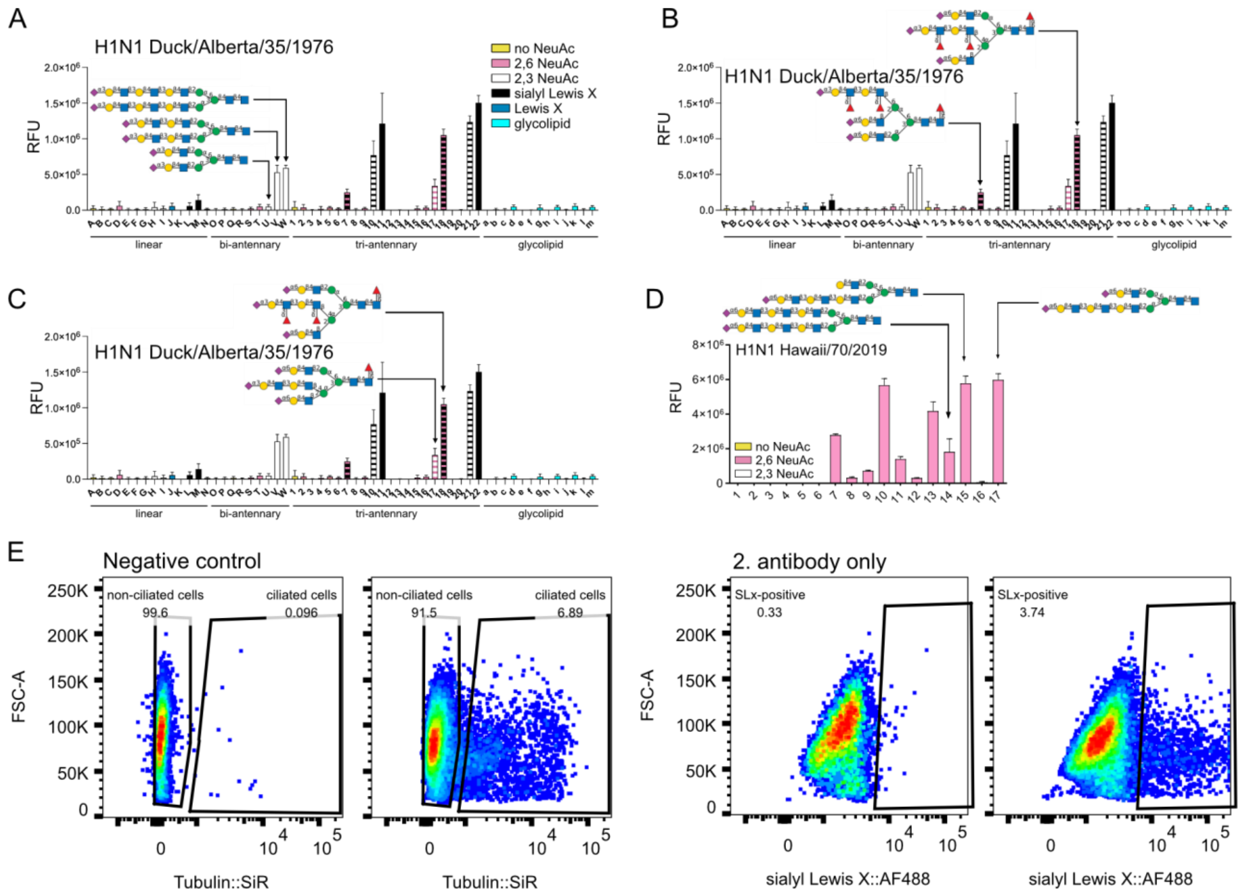


**Supp. figure 5. Avian H1N1 preference for sialyl Lewis X correlates with glycan abundance on ciliated cells**

**A) – C)** Microarray binding of H1N1 Duck/Alberta/35/1976 with glycans of Fig 4 A). The microarray tested for glycans with no sialylation (yellow), α2,6-linked NeuAc (pink), α2,3-linked NeuAc (white), Lewis X (blue), sialyl Lewis X (dark blue), and glycolipids (cyan). Striped bars indicate glycans terminating in different epitopes on different arms. Arrows highlight specific results. **D)** Microarray binding of H1N1 Hawaii/70/2019 with glycans of Fig 4 F). The microarray tested for glycans with no sialylation (yellow), α2,6-linked NeuAc (pink), and α2,3-linked NeuAc (white). **E)** Representative dot plots of gating strategy to determine non-/ciliated cells and apically sialyl Lewis X positive cells for Fig 5 D). BEpC from donor 2 were stained before trypsinization for apical sialyl Lewis X and ciliated cells (microtubule dye). The cells were analyzed by flow cytometry, cellular doublets were excluded by size, and dead cells with a cell viability marker.

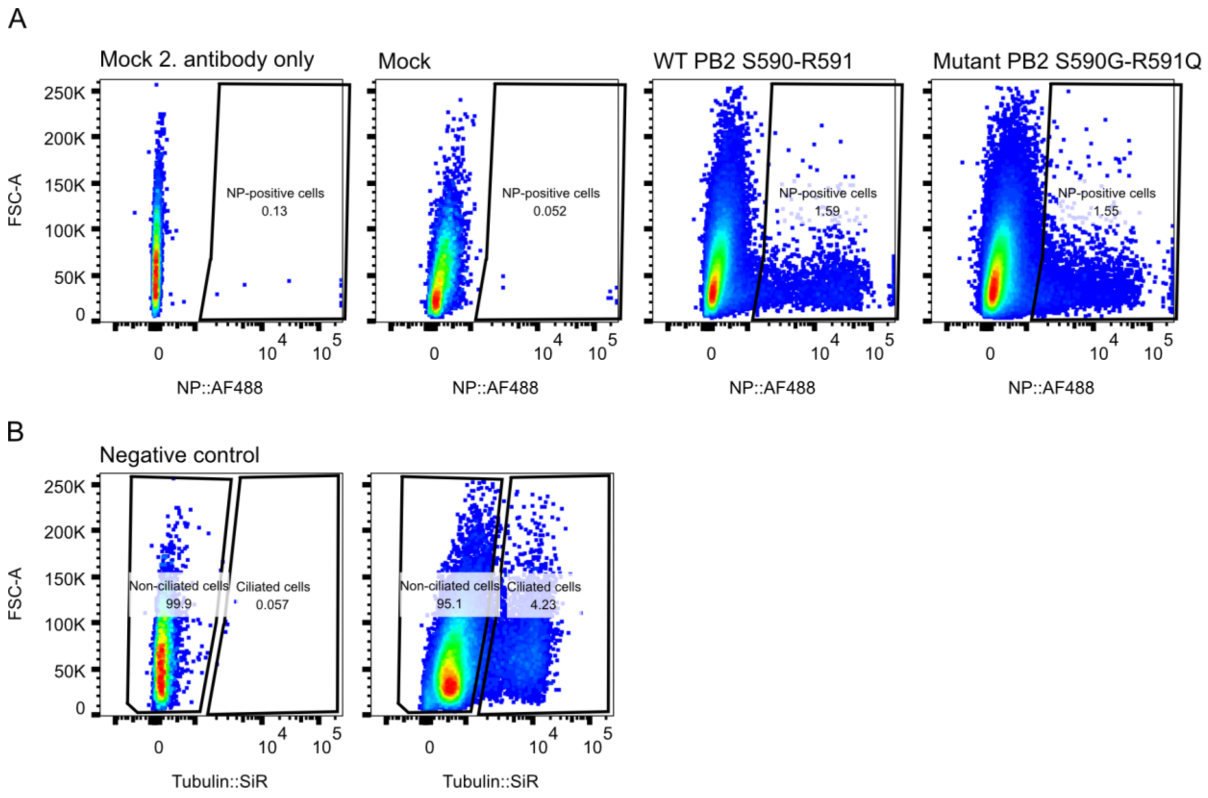


**Supp. figure 6. Early viral RNA and protein levels of human H1N1 are cell subtype specific**

**A) – B)** Representative dot plots of the gating strategy of A) NP-positive cells and B) ciliated cells for Fig 6 I). BEpC from donor 2 were infected with Netherlands/602/2009 PB2 S590-R591 or with Netherlands/602/2009 PB2 S590G-R591Q (MOI of 0.45). At 8hpi, cells were prepared for flow cytometry and stained for infected cells (NP-positive cells) and ciliated cells (microtubule dye). The cells were analyzed by flow cytometry, cellular doublets were excluded by size, and dead cells with a cell viability marker.
